# CascadeMAP: Autonomous Closed-loop Optimization of Enzyme Cascades via Microfluidics, Machine Learning and Agentic AI

**DOI:** 10.64898/2026.06.04.730034

**Authors:** Michal Vasina, David Kovar, Martin Kizovsky, David Lacko, Pavel Vanacek, Maximilian Herich, Eduard Volf, Lukas Drdla, Sona Cabalova, Pavlina Sikorova, Michael Jirasek, Pavel Solansky, Jan Jezek, Ota Samek, Filip Dousek, Hynek Walner, Pavel Zemanek, Andrew deMello, Zdenek Pilat, Jiri Damborsky, Stavros Stavrakis, Stanislav Mazurenko, Zbynek Prokop

## Abstract

Enzyme cascades enable complex biochemical transformations, but optimizing them requires searching high-dimensional spaces of enzyme stoichiometry, reaction conditions, and buffer composition. Here we introduce CascadeMAP, an autonomous microfluidic platform for closed-loop optimization of enzyme cascades, integrating high-throughput microfluidics with Bayesian optimization and assisted by agentic analyses. We demonstrate the platform across two three-enzyme cascades: (i) a glycerol detection pathway monitored by fluorescence and (ii) a 1,2,3-trichloropropane degradation pathway monitored by label-free Raman spectroscopy, providing orthogonal detection modalities. Bayesian optimization reached comparable optima with threefold fewer experimental conditions than Design of Experiments. A multi-agent AI system automated processing 2.1 GB of experimental data, pattern recognition, and insight synthesis. Operating without human intervention for 7 days, CascadeMAP processed >295,000 reactions across ∼7,400 different conditions. This capability establishes a generalizable, resource-efficient framework for autonomously optimizing enzyme cascades and metabolic pathways and accelerates the development of biocatalytic and synthetic biological systems.

## Introduction

Nature has composed an immense “songbook” of biotransformations, conducted by enzymes thanks to their high selectivity under mild conditions, which is exploited industrially in food, textile, agricultural, chemical and pharmaceutical processes.^1^ Traditionally, these processes relied on whole-cell metabolic engineering. However, cell-free enzyme cascades are increasingly utilized as a robust alternative, offering superior flexibility and fine-grained control over reaction variables.^2,3^ Despite the reduced biological complexity of cell-free systems compared to intact cells, the optimization of these cascades remains a significant challenge. Developing efficient biocatalytic systems requires the collection of high-fidelity datasets that capture the intricate interplay of multiple physicochemical parameters.^4^ This creates two central technical bottlenecks: (i) the high-throughput generation of high-quality multidimensional datasets, and (ii) the reliable analysis of these datasets to extract actionable insights and identify optimal operating conditions.

Currently, the leading technologies to tackle these challenges are microfluidic platforms, which enable controlled and high-throughput experimentation, and machine learning approaches (ML), recently enhanced by multi-agent AI research assistants,^5^ which ensure efficient navigation through complex multidimensional parameter spaces.^6^ Microfluidic systems have been applied to enzyme cascades, but their use has largely focused on proof-of-concept demonstrations, particularly involving enzyme immobilization and compartmentalization strategies, rather than the systematic, multidimensional optimization of cascade performance.^7^ In parallel, ML-based algorithms have demonstrated considerable potential for multiparameter optimization of complex reaction systems, including enzyme cascades.^8^ For example, the active learning-based software METIS has been successfully applied to optimize the CETCH cycle,^9^ highlighting the ability of ML to accelerate optimization of complex metabolic networks. Despite these advances, experimental implementation of ML-guided optimization has remained largely constrained by the throughput, reproducibility, and autonomy of the robotic liquid handling platforms.^10^ To date, integration of ML with microfluidic experimentation for enzyme cascades has been limited. Microfluidic continuous stirred-tank reactors (CSTR) have been coupled to active learning to infer kinetic models of enzymatic networks.^11,12^ That work was designed for model inference rather than closed-loop optimization across multidimensional reaction spaces.^13^ Moreover, despite the iterative nature demonstrated by METIS or CSTR, these systems do not run fully autonomously and require a human-in-a-loop for sample handling, reagent replenishment, or data transfer.^14^

Here we present CascadeMAP, a CascadeMAP, a microfluidic platform that combines classical design-of-experiments (DoE) with machine learning–guided optimization in a fully closed-loop workflow for enzyme cascade engineering.^15–17^ CascadeMAP enables the simultaneous, multidimensional optimization of key parameters critical for enzyme cascade development, including enzyme stoichiometry, reaction temperature, and pH, while autonomously executing experiments, acquiring data, and guiding subsequent experimental decisions. We validated CascadeMAP using two model systems, a three-enzyme glycerol detection pathway^18^ and a three-enzyme synthetic metabolic pathway for degradation of the industrial and environmental pollutant 1,2,3-trichloropropane.^19–21^ The platform replaces an operator-paced, reagent-limited screening campaign with an unattended search that consumes ∼0.2 µg of total enzyme and ∼1 min of instrument time per condition, and it identifies optimal operating conditions in multienzyme networks relevant to biocatalysis, biosensing and metabolic engineering.

## Results

### CascadeMAP hardware and software development

We developed CascadeMAP, an autonomous microfluidic platform for the optimization of enzyme cascade systems. The platform integrates multiple orthogonal detection modalities, including fluorescence-based assays and label-free Raman spectroscopy, enabling quantitative monitoring of reaction progress and intermediate formation. CascadeMAP supports reaction times ranging from minutes to hours and allows systematic exploration of multidimensional reaction spaces. The system operates in both conventional design-of-experiments (DoE) mode and a fully autonomous active learning (AL) mode, enabling efficient, data-driven optimization of complex enzymatic cascades (**Figure 1**).

**Figure 1.**
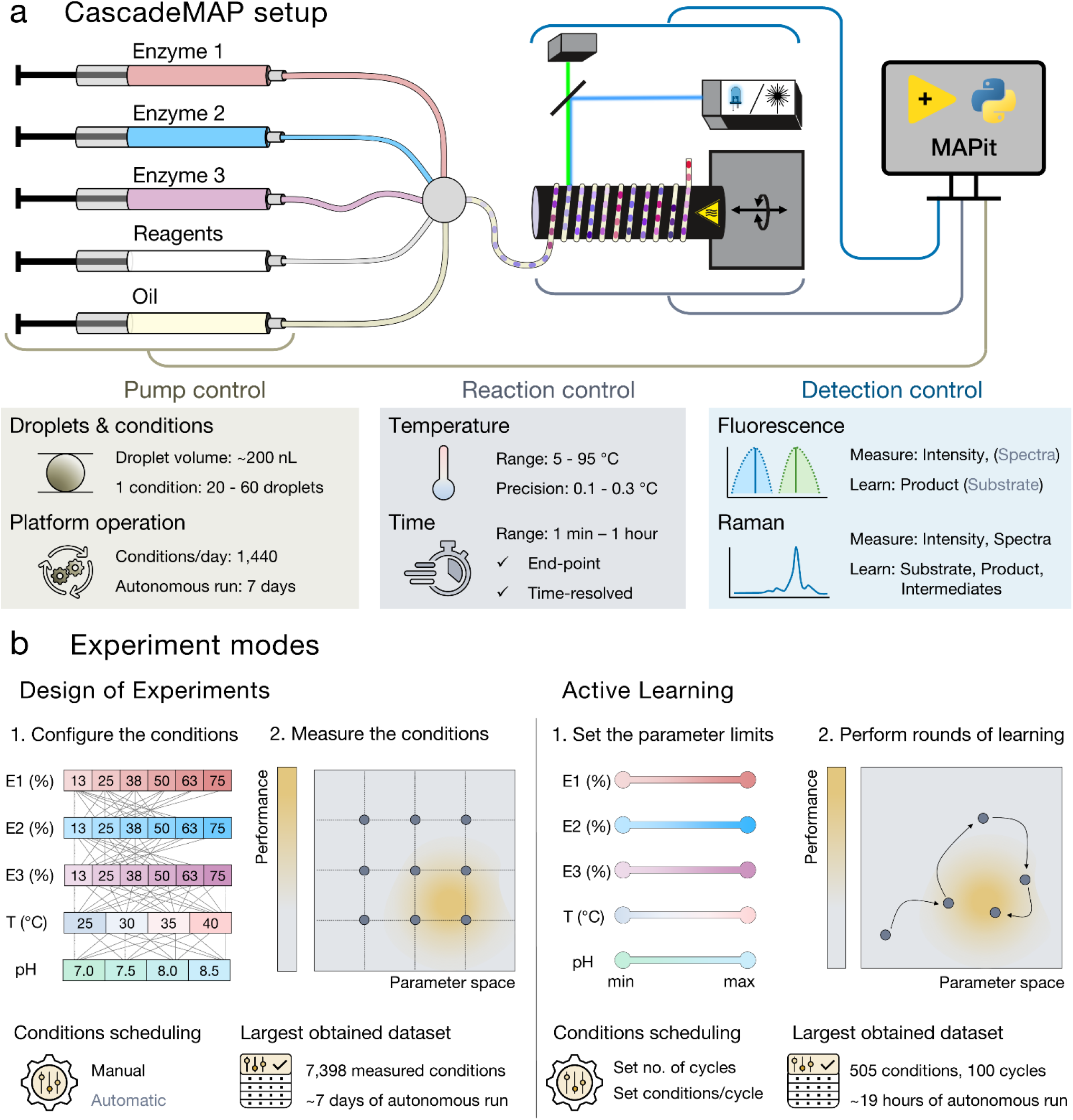
CascadeMAP microfluidic platform for closed-loop optimization of enzyme cascades. **a**, The technical setup of the microfluidic platform. The pumps control the flow rates of aqueous and oil phases, which are mixed in the manifold (grey circle), where droplets are also generated. The droplets travel through the incubation (reaction) coil wrapped around a temperature-controlled rod, where fluorescence can be read out in every loop of the tubing for time-resolved detection (Raman detection requires platform setup adjustments, see **Figure S12**). The software MAPit controls (i) the flow rates on syringe pumps (beige lines), dictating the experimental mode (see **b**), (ii) the reaction conditions, including temperature and reaction time (grey lines), and (iii) the acquisition of detected signal (blue lines). Details on control of pumps, reaction and detection are described below the platform scheme in beige, grey and blue squares, respectively. **b**, The experimental modes showcasing design of experiments (left) and active learning (right). Both modes are depicted as two-step processes of (i) defining the combinatorial space of conditions and (ii) their measurement. Typical parameters are enzyme concentrations (E1, E2, and E3), temperature (T), and pH.

During the development, we addressed several technical challenges inherent to conventional capillary-based microfluidic architectures. In typical designs reported in the literature,^22–25^ aqueous reagents are first mixed in a manifold, and droplets are subsequently generated at a downstream T-junction. The resulting axial dispersion between mixing and droplet formation smears concentration gradients, compromising temporal precision and forcing long delays between reaction conditions. CascadeMAP instead integrates the aqueous and oil phases in a single manifold, coupling mixing directly to droplet generation; the reduced dead volume suppresses dispersion and allows rapid, well-defined switching between flow-rate combinations (**Figure S1, S2**), which is difficult to achieve in conventional setups.

We implemented various technical solutions to preserve sample integrity and maintain stable experimental conditions during extended self-driving experiments, typically lasting more than 10 hours (**Figure S3**). A refrigerated enclosure preserved enzyme stability and gave a consistent signal over time at the cost of a larger footprint (**Figure S3**). To accommodate reactions spanning longer timescales and reach control of temperatures below room temperature, we also integrated a flask-based incubator and a complementary detection cell for end-point measurements. These additions extended the versatility of CascadeMAP, allowing it to monitor both rapid and slow enzymatic processes with equal precision (**Figure S4**).

Alongside the hardware development and optimization, we developed the MAPit software suite to automatically operate CascadeMAP and process the obtained data. The software architecture integrates a LabVIEW-based hardware control layer (National Instruments, US) with Python-based routines responsible for inline data processing and execution of the active learning algorithm. The system enables fully autonomous execution of both DoE and machine learning–driven experiments, including extended runs of up to 7 days, with robust, unattended operation (**Figure S5**). In a DoE robustness test, the platform autonomously generated and analyzed >295,000 reactions across 18 temperature set points from 20 to 60 °C, establishing fluidic and optical stability over this interval (**Figure S5**).

### Design of Experiments on CascadeMAP

#### DoE optimization of the glycerol detection cascade

As a model system, we selected the glycerol detection pathway (**Figure 2**), adapted from the method reported by Morita and Terada.^18^ This cascade involves three enzymes, glycerol kinase (GK), glycerophosphate oxidase (GPO), and horseradish peroxidase (HRP), to generate a fluorescence-based output, well-suited for the validation of the CascadeMAP architecture. The fluorophore used in the original method, Amplex Ultra-Red (AUR), performed satisfactorily on a microtiter plate (MTP) reader but gave irreproducible signals and compromised droplet stability on CascadeMAP due to precipitation, as reported previously by Zhu et al.^26^ (**Figure S6**). We therefore transitioned to aminophenyl fluorescein (APF), a fluorogenic probe that becomes fluorescent upon reaction with highly reactive oxygen species and is employed in various assays.^27,28^ The APF was successfully validated for performance on both the MTP reader (**Figure S7**) and CascadeMAP, providing the stability and signal reproducibility required for all subsequent autonomous experiments.

**Figure 2.**
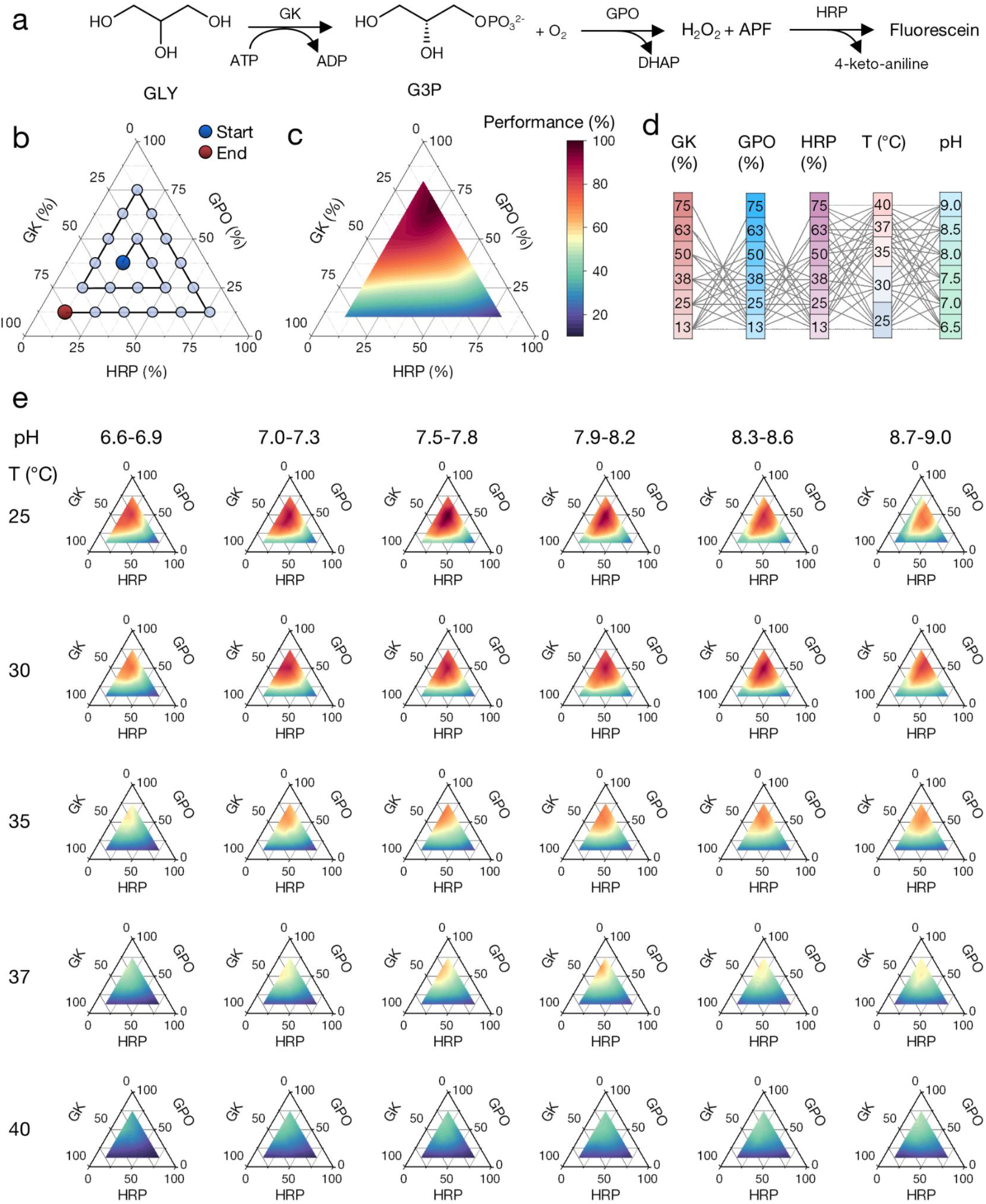
The DoE optimization of the glycerol detection pathway. **a**, General scheme of glycerol detection pathway. Glycerol kinase (GK) converts glycerol (GLY) to glycerol-3-phosphate (G3P), while adenosine triphosphate (ATP) is dephosphorylated to adenosine diphosphate (ADP). G3P further reacts with oxygen, yielding dihydroxyacetone phosphate (DHAP) and hydrogen peroxide, which, catalyzed by horseradish peroxidase (HRP), oxidizes aminophenyl fluorescein (APF) to form well-detectable fluorescein. **b**, The design of the experiment is based on a ternary plot diagram, where the axes represent enzyme ratios. The graph also shows the sequence of measurements from the beginning (blue point in the middle) to the end (red point in the bottom left corner). **c**, The resulting ternary plot from enzyme ratios optimization at 37 °C (30 droplet per condition). The colormap shows the relative fluorescence signal translated to performance of the cascade in %. **d**, The parameter space and combinations of conditions for 5-parameter DoE (enzyme ratios, temperature, and pH). **e**, Ternary plots of 5-parameter DoE for 6 different pH values (columns, expressed as ranges due to variation with temperature) and for 5 different temperatures (rows). The colormap shows the relative fluorescence signal in %, the same as in **c**.

Our initial objective was to optimize the relative ratios of enzymes within the glycerol detection cascade while maintaining a constant total enzyme concentration (**Figure 2b**). First, we chose a “safe time-resolved detection” mode enabling time-resolved detection across multiple loops on the rod while preventing photobleaching of the APF fluorophore (**Figure S8b,c**). This mode was effective for kinetic characterization but detected only one half of the generated droplets, increasing both reagent consumption and run time (**Table S1**). Leveraging the improved droplet generation stability of CascadeMAP (**Figure S2**), we tested a “stepwise end-point detection” mode (**Figure S9**) that maximized the throughput of detected droplets, significantly reducing sample requirements and total experimental time (**Figure 2c** and **Figure S8d, e**). More specifically, for 36 enzyme ratios, the step-wise end-point mode needed 0.66 h and 7.7 µg of enzyme, compared with 7.2 h and 83 µg in the safe time-resolved mode (about 11-fold less time and enzyme consumption; **Table S1**). Validation experiments confirmed that both detection modes yielded consistent results, identifying an optimal GK:GPO:HRP ratio of approximately 2:6:2 (**Figure S8b, e**).

To target the optimization of multiple parameters within a single experiment, we expanded our experimental scope to include reaction temperatures and pH, both of which are key parameters influencing the performance of enzyme cascades.^7^ Optimization of reaction temperature was facilitated by the temperature-controlled rod-shaped incubator as introduced previously.^24,25^ To vary the pH, we employed two separate syringes delivering 50 mM Tricine and 50 mM Bis-Tris propane, achieving a linear pH range from 6.5 to 9.5 and capturing both the upper and lower activity limits of the cascade (**Figure S10**). We abandoned in-line pH monitoring because the available flow cell introduced unacceptable dispersion (**Figure S11**). Instead, pH was calibrated off-line at each reaction temperature (**Figure S10, Table S2**).

Building upon the initial enzyme ratio screenings, we expanded the DoE to combinatorially explore five parameters: 22 different enzyme ratios, 5 different temperatures, and 6 different pH levels - in total 660 unique reaction conditions with ∼20 droplets per condition (**Figure 2d**). This exhaustive screening was completed in approximately 12 hours using endpoint measurements, identifying the optimum at 25 °C and pH 7.6 (**Figure 2e**).

#### DoE optimization of the TCP degradation pathway

To expand the application scope of CascadeMAP, we combined the glycerol detection pathway with a previously kinetically characterized synthetic metabolic pathway for degradation of 1,2,3-trichloropropane (TCP pathway) (**Figure 3a**).^19^ To accommodate the longer reaction times from 5 to approximately 30 min, required for TCP conversion to glycerol, we implemented a heated reaction zone with an end-point detection cell (**Figure S4**). Optimization of the TCP pathway enzyme ratios (**Figure 3c**) yielded results consistent with previously reported optimal ratios (**Figure 3b**).^19^

**Figure 3.**
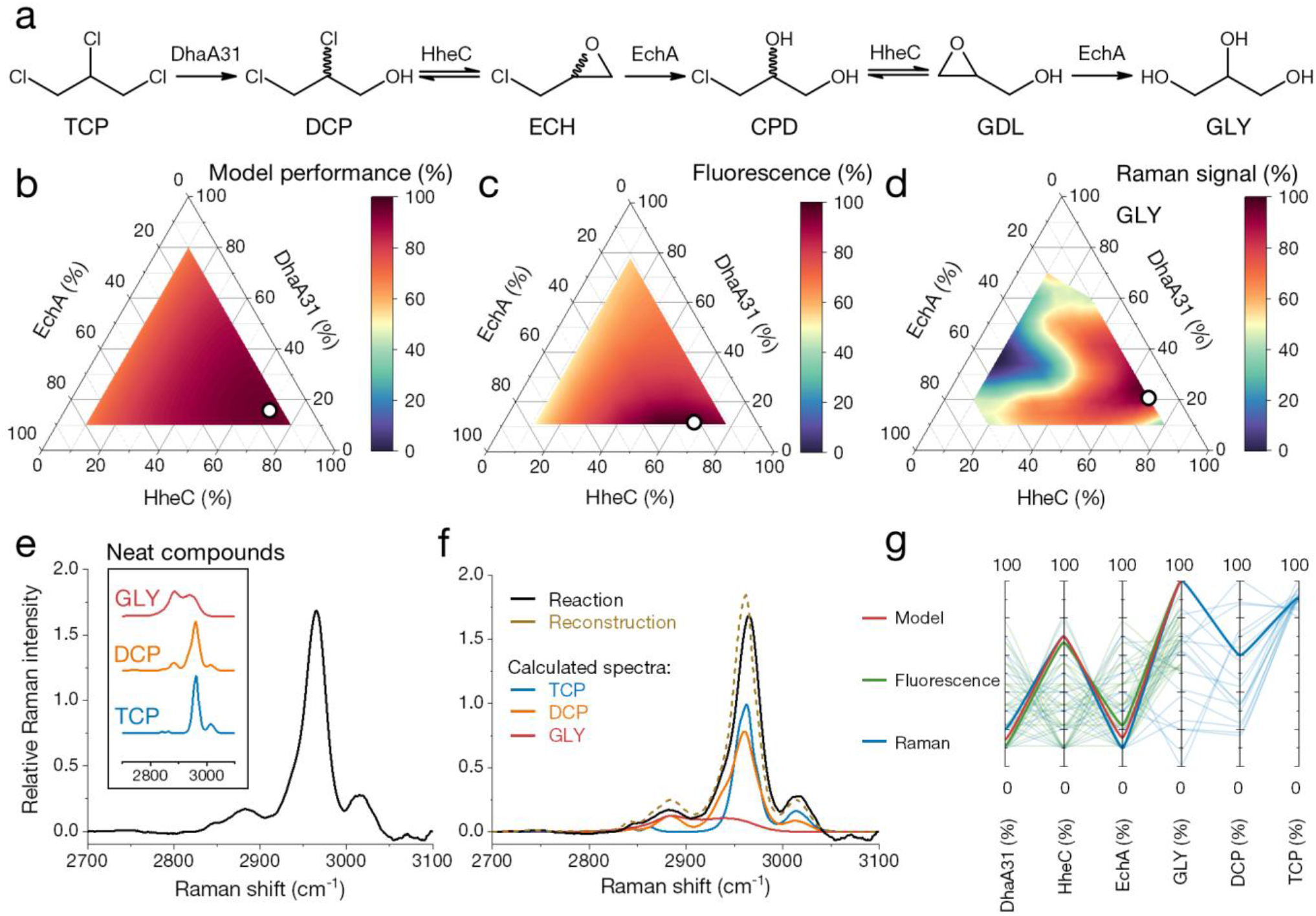
Optimization of enzyme ratios for TCP degradation pathway. a, The scheme of the TCP degradation pathway. The abbreviated compound names denote: TCP – 1,2,3-trichloropropane, DCP – (*R,S*)-2,3-dichloropropane-1-ol, ECH – (*R,S*)-epichlorohydrin, CPD – (*R,S*)-3-chloropropane-1,2-diol, GDL – (*R,S*)-glycidol, and GLY – glycerol. **b**, The prediction of TCP pathway performance depending on enzyme ratios based on a kinetic model reported previously.^19^ **c**, Relative fluorescence signal of TCP pathway enzyme ratio optimization employed the glycerol detection pathway. The measurement was conducted at ∼30 min of the reaction time, and the mean reaction temperature was 38.2 ± 0.2°C. **d**, Relative Raman signal depending on enzyme ratios for the product. The experiment was carried out at ∼30 min of reaction time, and the mean reaction temperature was ∼37°C. **e**, The Raman spectrum for the enzyme ratios producing the most GLY with spectra of the neat compounds (inset). **f**, The overlay of the reaction spectrum (black) from **e** and the spectrum (beige) reconstructed from non-negative least squares (NNLS) calculation of individual compounds. **g**, The parallel coordinates plot of combined datasets from fluorescence (**c**, green lines) and Raman (**d**, blue lines) and the top ratio combination from the kinetic model (**b**, red line). Each dataset was normalized to the maximum value of that dataset. These best conditions (i.e., those showing maximum glycerol production) are highlighted with bold lines and correspond to white circles in **b-d**. The lighter lines denote remaining conditions (one line for every condition). DCP and TCP axes are shown only for the Raman dataset, as for the other datasets, these values were not measured or predicted.

To demonstrate the versatility of CascadeMAP beyond fluorescence-based detection, we adapted the system to label-free monitoring of the TCP pathway using Raman spectroscopy (**Figure S12**). This required integrating a glass capillary into the setup to define a stable detection zone (**Figure S12b**), developing a specialized capillary–tubing interface that preserved droplet monodispersity (**Figure S12c**), and implementing custom software for automated real-time spectral acquisition and processing. To enable the continuous detection of low concentrations of metabolites in flowing droplets, we employed a high-power Raman microspectrometer developed at the Czech Academy of Science (ISI CAS, Brno, Czech Republic), featuring a 6 W laser. To our knowledge, this high-power approach has not previously been applied to enhance Raman sensitivity in microfluidic droplet systems, enabling the analysis of metabolites at concentrations below the limit of detection of conventional Raman spectrometers.

By leveraging the label-free detection capabilities of Raman spectroscopy, we successfully detected the substrate (TCP), the first intermediate (*R,S*)-2,3-dichloropropane-1-ol (DCP), and the product glycerol (GLY) in real-time. While other intermediates (**Figure 3a**) were also detectable as neat compounds (**Figure 3e,f, Figure S13, Table S3**) their concentration in the reaction mixture was not accounted for as preliminary results indicated that they do not accumulate during the reaction. Thus, the analysis was solely focused on TCP, DCP, and GLY relative contributions to the resulting spectra (**Figure S14** and **S15, Table S3**). The optimization of enzyme stoichiometry using Raman-based detection (**Figure 3d, Figure S14** and **S15**) mirrored the patterns observed in our fluorescence data (**Figure 3c**) and fitted well with established kinetic model predictions (**Figure 3b**).^19^ Notably, comparing the top-performing enzyme ratios showed that both the fluorescence- and Raman-based measurements identified an optimum consistent with the kinetic model (**Figure 3g**).^19^ Furthermore, CascadeMAP allowed us to identify the range of enzyme ratios that lead to DCP accumulation (**Figure S14c**). TCP concentration remained approximately constant due to continuous oil feeding (**Figure S14b**), falling slightly where DCP accumulated, suggesting slow substrate depletion within the droplets, and rising where DhaA and HheC were both too dilute to initiate degradation.

#### Bayesian optimization of the glycerol detection pathway

To fully harness the autonomous nature of CascadeMAP experiments, we incorporated a Bayesian optimization (BO) active learning (AL) module that operates in a closed-feedback loop even during experimental runs (**Figure 1b** and **S16**). The user defines the parameter ranges to be optimized and sets the “budget”, meaning the number of conditions to be tested in the initial and subsequent rounds (**Figure S17**). Then the module integrated in MAPit proceeds as follows. Initial conditions are drawn by quasi-random Halton sampling, while maintaining constant total flow of selected input, e.g., enzyme solutions.

To minimize stabilization delays, the algorithm also orders conditions by changing high-latency parameters, such as temperature, in one direction and solving a weighted Travelling Salesperson Problem to minimize changes in the remaining parameters between consecutive conditions.

The data generated in the initial round via the “step-wise end-point” acquisition mode are processed and analyzed by the MAPit software in quasi-real-time (**Figure S8d, S9** and **S18**) and fed into the BO algorithm, which creates the model of the reaction landscape and proposes a next batch of conditions to be tested (**Figure 4**). After measuring these conditions, the data are fed back to the BO algorithm to update the model. This entire process—measuring, data processing and analysis, learning, and re-planning—proceeds iteratively in a closed loop, enabling fully autonomous operation for days at a time without any human intervention.

**Figure 4.**
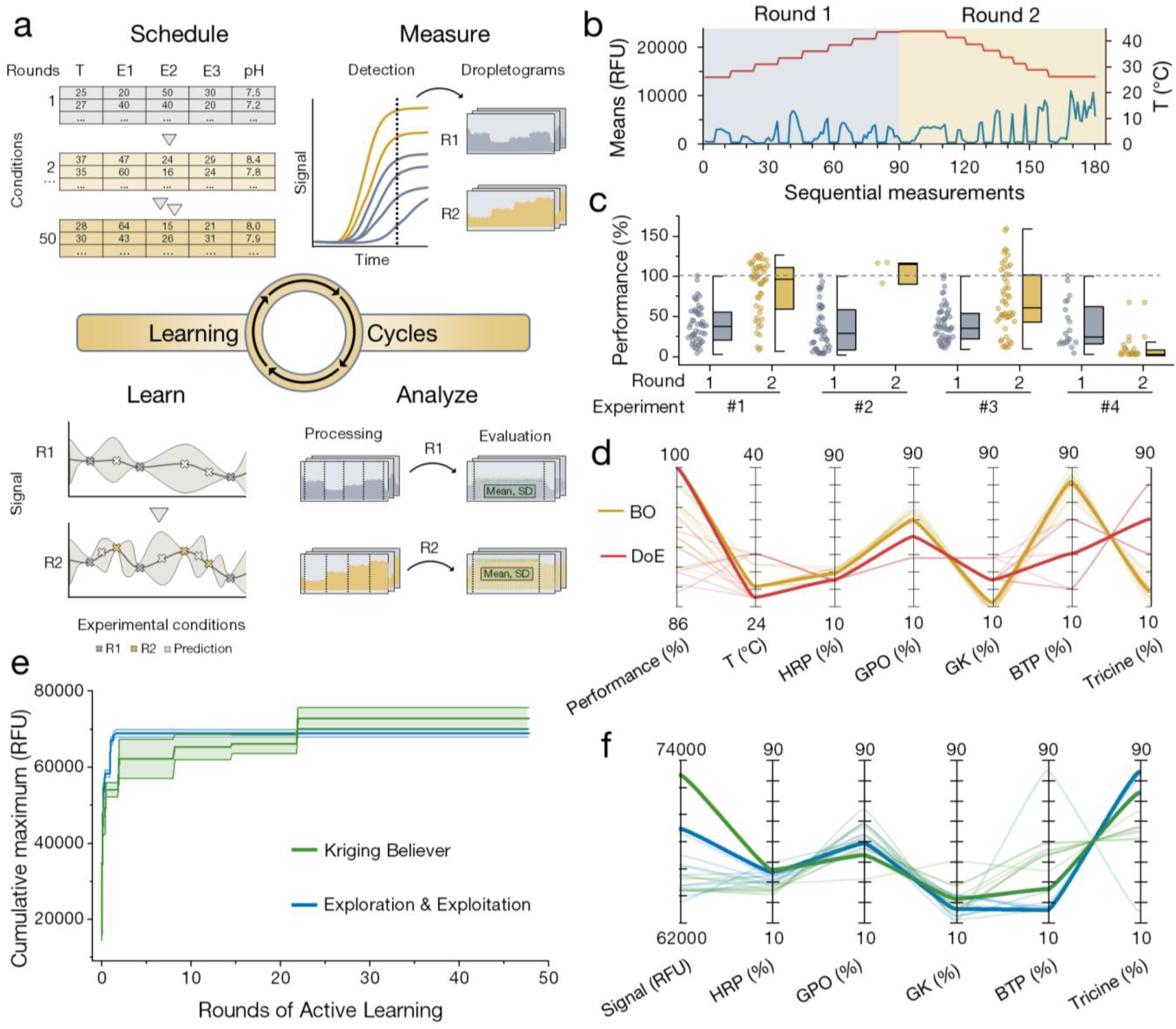
Active learning optimization of the glycerol detection pathway. **a**, Workflow of active learning (AL) involving (i) scheduling the conditions to test in each round based on the limits for each parameter to be tested, (ii) measurement of data involving end-point detection, (iii) processing and evaluation of data, and (iv) learning of patterns in data and updating the model to predict new conditions (white crosses). **b-c**, AL optimization of temperature (T), pH, and enzyme ratios. **b**, Example of data acquired during the initial and next round showing means of fluorescence signal for every condition (blue line, expressed in relative fluorescence units, RFU) and temperature (red line). **c**, Box plot summarizing four experiments composed of two rounds of AL. The performance on the Y axis denotes a signal relative to the maximum signal obtained during the first round of a particular experiment, as indicated also by the gray dashed line. Experiments 1-3 included 50 initial conditions to be measured, while experiment 4 had 20 initial conditions measured. For experiment 2, only 3 of the next conditions were measured. **a-c**, Rounds 1 (R1) and 2 (R2) stand for the initial and the subsequent rounds of AL and are marked by grey and yellow, respectively. **d**, Parallel coordinates plot comparing the 10 conditions with the highest signal from DoE (red, Figure 2e) and AL (gold, **c**) when optimizing enzyme ratios (horseradish peroxidase (HRP), glycerophosphate oxidase (GPO), and glycerol kinase (GK)), temperature, and pH (ratio of Tricine and Bis-tris propane (BTP)). The best conditions from each algorithm are connected by a bold line. **e-f**, AL optimization of pH and enzyme ratios using two AL algorithms, Exploration and Exploitation (blue) and Kriging Believer (green). The data from the KB dataset were linearly projected onto the EE dataset (**Figure S20**). **e**, The cumulative maximum of observed signal throughout 48 rounds of active learning. The lines show means of the condition with maximum signal, and the shaded band shows the standard deviation for the particular condition. **f**, Parallel coordinates plot of the 10 highest-signal conditions from each algorithm. The best conditions from each algorithm are connected by a bold line (**d, f**).

We validated the AL workflow using the glycerol detection pathway as a straightforward model cascade with relatively short reaction times. We sought to optimize enzyme ratios, pH, and temperature. The first challenge was to automate the BO algorithm’s learning from the initial set of measured conditions. We performed several optimizations with a single next round of active learning (**Figure 4b-c**). Our results demonstrate that the BO algorithm successfully identified higher-performing conditions in the second round, consistently outperforming the best results from the initial sampling (**Figure 4c**). However, we noted a critical dependency on the experimental “budget”: when the number of conditions per round was too low relative to the complexity of the experiment (e.g., **Figure 4c**, Experiment 4 with 20 conditions per round), the overall combinatorial space was undersampled, and the second round failed to yield a substantial enrichment. When we compared the relative performances of the previous DoE optimization and the current single-next round AL dataset, we found the identified optimal temperature and enzyme ratios to be highly consistent (**Figure 4d**). Despite a differently identified pH optimum, the AL algorithm could efficiently sample the combinatorial space (**Figure S19**) while providing optimum using only one third of conditions (225 conditions) compared to the DoE optimization (660 conditions, **Figure 2e**), thus substantially increasing experimental efficiency.

To evaluate the robustness of the AL workflow over extended durations, we transitioned from single-round optimization to a multi-round sequence at a fixed reaction temperature, with 10 conditions in the initial round and 5 conditions per subsequent iteration. During these extended runs, we compared two different batch-sequential strategies: the formerly used Exploration and Exploitation (EE) algorithm^29^ balancing exploration and exploitation, and the Kriging Believer (KB) algorithm^30^ guiding the optimization more towards exploitation. We compared the first 48 rounds of AL (**Figure 4e**), corresponding to 245 autonomously measured conditions. Due to minor mechanical misalignments between the temperature-controlled incubator and the optical detection system, the raw datasets were not directly comparable across different runs. Because the initial 10-condition sequences were identical for every experiment, we used these points as an internal reference to project the KB dataset linearly onto the EE one (**Figure S20**). Both strategies identified similar optima; the primary divergence appearing in the pH buffer ratios, where the KB algorithm slightly outperformed the EE approach (**Figure 4e-f**).

#### Analysis of Bayesian optimization datasets by agentic AI systems

The concept of autonomous self-driving laboratories aligns with the emerging paradigm of multi-agent AI systems.^5,31^ We explored the capabilities and limitations of the multi-agent AI system Theorema Lab 1.0.9^32^ on a 5-prompt task of autonomous dataset analysis covering peak processing, alignment of acquisition rows with conditions, multivariate statistics (principal component analysis and SHAP^33^), and interactive reporting (**Figure 5a**). These prompts were executed on the two representative BO datasets (EE and KB, **Figure 4e-f**) by using the harness engineering approach.^34^ The performance of Theorema Lab 1.0.9 was compared to Claude Opus 5,^35^ Deepseek V4 Flash, ^36^ GLM 5.2^37^ and Kimi K3.^38^ Unlike the other systems (**Figure S21**), the runs using GLM 5.2 and Kimi K3 failed the first prompt, where means of processed peaks were checked against the automated peak analysis by weighted Deming regression (accepted slope within 5% of 1.0, **Figure S22**) and were therefore excluded from further testing. For the remaining prompts, Theorema Lab 1.0.9, Claude Opus 5, and Deepseek V4 Flash delivered the requested outputs with similar findings. Exploration curves and principal component analysis indicated that EE favored global exploration, whereas KB favored exploitation (**Figure 5b,c**). SHAP analysis identified higher GPO levels (40–60%) and intermediate HRP concentrations (25–40%) as favorable for the target response, consistent with GPO being rate-limiting (**Figure 2e, 4d,f**), while BTP, Tricine, and GK acted mainly as local modulators (**Figure 5f,g**).

**Figure 5.**
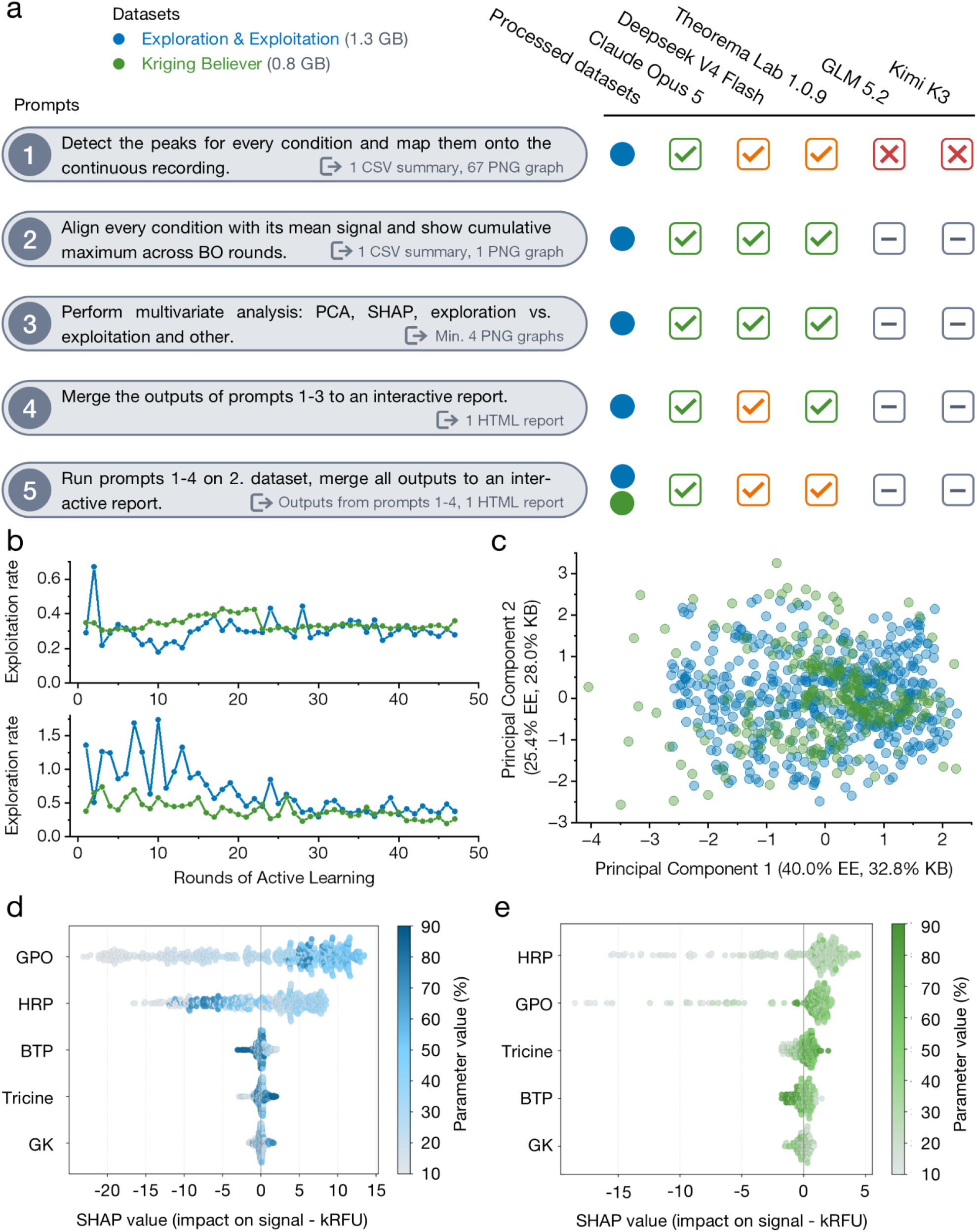
The analysis of BO data by agentic AI systems. a, The overview of analysis of BO datasets (Exploration & Exploitation blue and Kriging Believer green) analysis by means of agentic AI. The workflow is described by shortened prompts (left, grey ovals) with expected outputs (gray, bottom right). The high-level evaluation of how the particular AI system fulfilled the prompts is shown by checkboxes denoting satisfactory results with (orange checkbox) and without (green checkbox) any necessary human intervention, non-satisfactory results (red cross). Grey minus signs denote that the AI system was not prompted. The abbreviations include comma-separated values (CSV), portable network graphics (PNG), principal component analysis (PCA), SHapley Additive exPlanations (SHAP), and hypertext markup language (HTML). **b-e**, Selected outputs of the analysis performed by Theorema Lab 1.0.9. **b**, The progress of exploration (bottom) and exploitation (top) for both algorithms, where exploration measures how far a particular round moves from earlier measured conditions, while exploitation measures closeness to the condition with currently maximum observed signal. **c**, The score plot of principal component analysis with the percentages of variance that each principal component explains in the brackets of the particular axis label. **d-e** SHAP analysis^33^ assessing the influence of each parameter (enzymes and pH buffer percentage) on the resulting signal in thousands of relative fluorescence units (kRFU) of the Exploration & Exploitation (**d**) and Kriging Believer (**e**) dataset. Each point represents an experimental condition, positioned by SHAP value (impact on signal), while color intensity reflects the corresponding feature value.

Regarding the soft metrics of agentic analyses, Theorema Lab 1.0.9 used the lowest number of tokens and less than half the tool calls of other systems, but a five-hour stall made it the slowest, whereas Claude Opus 5 completed all tasks in approximately 2.5 hours without human intervention but consumed 6.5–7.5 times more tokens and was the most expensive (**Table S4**). Deepseek V4 Flash was the cheapest at $1.96, but required human corrections and left minor deliverable requirements unmet. Together, these results demonstrate that agentic AI systems can significantly augment the analysis stages of automated experimentation. Ongoing developments are focusing on expanding this framework to support on-the-fly learning experiment execution. However, a paramount emphasis remains on integrating rigorous human-in-the-loop control, operational guardrails, and strict safety protocols to govern autonomously operating physical systems.^5^

## Discussion

The multiple-parameter optimization of enzyme cascades provides clear benefits from industrial bioprocessing aimed at maximizing product yields to metabolic engineering focused on deciphering the intricate flux of synthetic networks. In this work, we introduced CascadeMAP as a platform that efficiently combines the high-throughput capabilities of droplet microfluidics^6^ with machine learning^4^ for exploring patterns in complex data, enabling accelerated data-driven optimization. The number of parameters to be simultaneously optimized on CascadeMAP is primarily determined by the number of inlet ports on the fluidic manifold. Commercially available manifolds offer up to 16 inlets, which theoretically enable up to 15 factors to be varied (accounting for oil required for droplet generation). Expansion beyond this limit requires custom manifolds or lab-on-a-chip devices, which could further facilitate high-resolution, time-resolved detection as previously described.^39,40^ External parameters such as temperature and reaction time are regulated within the same closed loop: CascadeMAP varied enzyme stoichiometry, temperature and pH in a single DoE campaign, and enzyme stoichiometry and pH over up to 100 consecutive active-learning rounds. Further versatility and control over sample quality can be achieved by integrating pressure-driven pumps^41^ and customized temperature control of samples to be introduced to CascadeMAP.

During CascadeMAP development, we found that selecting a suitable analytical assay is critical. While Amplex UltraRed^26^ works well in microtiter plate assays, its instability during the multi-hour storage required in syringes makes it incompatible with long-term autonomous runs. A switch to aminophenyl fluorescein (APF),^27,28^ which offers excellent stability, solved these issues, yet still allowed only monitoring of the end product. To extend the information content yielded from the reactions, we introduced high-power label-free Raman detection, enabling concurrent monitoring of substrates, intermediates, and products, exemplified by the degradation of the industrial pollutant TCP, for which a packed-bed biocatalytic process has been demonstrated.^20^ Such sensitivity exceeds conventional Raman systems and avoids the non-linear signal enhancements and complex nanoparticle substrate requirements of Surface-Enhanced Raman Spectroscopy (SERS), which complicate quantitative interpretation.^42^ Future versions of CascadeMAP may integrate absorbance, luminescence, or mass spectrometric detection for even greater versatility.

Recent advances in autonomous experimentation highlight how coupling machine learning with robotic systems can significantly accelerate biochemical research.^15,43,44^ Microfluidic devices such as CascadeMAP extend these gains further, offering high throughput at reduced reagent consumption together with precise temporal control and real-time adjustment of experimental conditions.^6^ Placed against other self-driving laboratories, CascadeMAP operates at a different scale. Campaigns on robotic platforms targeting biological optimizations were run in typical reaction volumes of 10-200 µL at tens to lower hundreds of measured conditions per day,^9,13,45–47^ whereas CascadeMAP measured conditions in droplets of 200-500 nL at ∼650 (active learning) to ∼1,300 (DoE) conditions per day. These enzyme platforms completed 4–38 closed-loop cycles; while CascadeMAP completed 100 fully autonomous active learning rounds.

A particularly promising direction within this field is the emergence of agentic AI systems.^5^ We assessed five agentic AI systems on the autonomous processing and analysis of 2.1 GB of raw data across five prompts spanning peak processing, visualization, multivariate statistics, and interactive reporting. These analyses could be finished within ∼3 hours and provided useful dataset-level insights, which would be tediously achieved by manual analysis (estimated at 85-115 hours of dedicated time). Future iterations of CascadeMAP will leverage agentic AI to plan and drive active learning experiments. By dynamically adjusting the balance between exploration and exploitation, the system will execute real-time, multi-objective optimization across coupled parameters, including product yield, selectivity of enzymatic reactions, suppressed inhibitions, and long-term operational stability, i.e., areas where agent-based Bayesian strategies have already shown effectiveness.^48,49^

In summary, CascadeMAP represents a robust platform for optimizing complex enzyme cascades within a single, unattended experimental campaign. Thanks to a high degree of fluidic automation and integration of machine learning, tens to hundreds of consecutive learning cycles can be carried out without any human intervention. By autonomously conducting experiments, CascadeMAP will contribute to the generation of high-quality datasets, requested by current ML pipelines,^50^ and will become a valuable tool for metabolic engineers to optimize complex pathways leading to value-added compounds.^51^

## Online methods

### CascadeMAP platform and experiments

#### Hardware

The CascadeMAP hardware is based on the previously introduced and thoroughly described KinMAP.^23^ Briefly, oil and aqueous phases held in GASTIGHT syringes (Hamilton, USA) are driven by low-pressure neMesys pumps (Cetoni, Germany) through high-purity PFA tubing to a 6-inlet (P-170, IDEX Health & Science, USA) or 10-inlet (Z10M1PK, Vici Valco, France) manifold, in which mixing and droplet generation are coupled within a single element. The droplets then travel through a reaction coil wound around a copper rod that is heated by an embedded cartridge and regulated by a PID controller from ambient temperature up to 180 °C, with an observed variation from the set point of 0.1 °C, and fluorescence is read out loop by loop. The ratios of solutions and the velocity of droplet movement were controlled by changing the flow rates of the individual pumps, keeping constant total flow rates of 12 and 12 μL/min for the aqueous and oil phases, respectively. Long-term experiments used automated refilling of the aqueous solutions through 3/2-way valves (Cetoni, Germany) and a dual-syringe oil supply unified by a Y-junction. A complete list of components, the optical configurations used for the APF and AUR fluorescence assays, and the extensions for sub-ambient and long-reaction-time operation are given in **CascadeMAP experiments with fluorescence detection** section of **Supplementary Information**.

#### MAPit software and experimental modes

The MAPit software was written in LabVIEW (National Instruments, USA) and is based on a queued message handler, which facilitates multiple sections of code running in parallel and enables control of the MAP hardware components. The main code was extended by Python-based scripts for accelerated peak detection and evaluation of a sequence of measurements. MAPit provides three experimental modes – sequence, flow profile and manual operation – and two data acquisition modes (**Figure S9**). The dispersion-tolerant “safe time-resolved” mode, inherited from KinMAP, first fills the reaction zone with droplets of the desired composition and only afterwards detects the signal in the loops of choice in a backward direction. The suppressed dispersion of the integrated manifold additionally enabled a “step-wise end-point” mode, in which a reaction condition is generated and detected within the same time interval (typically 1 min) and the signals are subsequently realigned to the correct conditions using auxiliary rows appended to each condition sequence. All modes are described in detail in **Hardware** section of **Supplementary Information**.

#### Bayesian optimization module

Additionally, a Bayesian optimization (BO) feedback loop was integrated into MAPit, providing online replanning of experiments based on the data obtained in the previous round. The BO module was written in Python and is available on GitHub: https://github.com/loschmidt/CascadeMAP-Bayesian. Briefly, the initial and subsequent sets of parameters were generated from quasi-random Halton sequences using the *scipy.stats.qmc.Halton* class of the SciPy library v1.17.0, with simplex-constrained parameter groups transformed according to the “transformation root” method^52^ and the execution order optimized by solving a weighted Travelling Salesperson Problem. The BO was based on Gaussian Process Regression implemented using the scikit-learn library v1.8.0, with expected improvement used as the default acquisition function. Two algorithms were implemented to account for batches at each iteration: Exploration-Exploitation sampling of different weights in the Weighted Expected Improvement function^29^ and Kriging Believer.^30^ Full implementation details, including the user-tunable exploration bias and the handling of duplicate suggestions, are given in the **Bayesian optimization scripts** section of **Supplementary Information.**

#### Reaction mixtures and conditions

In the glycerol pathway experiments, individual enzymes were loaded in separate syringes at concentrations such that the final reaction concentration of all enzymes summed to 1-3 U/mL, while glycerol, APF and ATP were mixed in one syringe so that the final reaction mixture contained 250 μM glycerol, 25 μM APF and 2 mM ATP; specific experiments required these concentrations to be adjusted accordingly. For the TCP pathway, substrate delivery was achieved through substrate partitioning between the oil (continuous phase) and aqueous (dispersed) phase as described previously.^23,53^ The enzymes were loaded either mixed or separately in syringes with a final reaction mixture concentration of 0.6 mg/mL, and the reaction time was set to about 30 minutes. TCP was supplied from fluorinated oil FC-40 (3M, USA) containing 0.25 % AZ2C surfactant (CF Plus Chemicals, Czech Republic, manufactured according to a previously published method^54^), at ∼9.4 mM for fluorescence and ∼47 mM for Raman detection. Buffer compositions, oil/surfactant preparation and the off-line pH calibration procedure are detailed in **Reaction mixtures and off-line calibration** section of **Supplementary Information**. Reference microtiter plate measurements were performed on the CLARIOstar plus reader (BMG Labtech, Germany) using 96-well plates with 100 μL reaction volume; the excitation and emission settings for AUR and APF are given in **Microtiter plate reader experiments** section of **Supplementary Information**.

#### Preparation of enzyme samples

Glycerol kinase from *Cellulomonas* sp. (G6142), Glycerol 3-phosphate oxidase from *Pediococcus* sp. (G9637), and Horseradish peroxidase (SRE0082) were purchased from Merck, USA. Upon dissolution, the enzyme concentrations were adjusted to 40 – 100 U/mL, flash-frozen in liquid nitrogen, and stored at -70 °C. For CascadeMAP measurements, they were transferred to 50 mM Tris-SO_4_ buffer, pH 8.0. Genes encoding DhaA31, EchA wt, and HheC wt were cloned into pET vectors and expressed in *E. coli* BL21 (DE3) cells. DhaA31 and EchA were purified from cell lysates by immobilized metallo-affinity chromatography, while HheC wt was purified using anion-exchange chromatography^55^ followed, for part of the fractions, by size exclusion chromatography. Plasmids, chromatographic gradients, buffer compositions and storage conditions are detailed in **Preparation of enzyme samples section** of **Supplementary Information**.

#### Raman detection

Label-free detection used an in-house-built Raman microspectrometer developed at the Institute of Scientific Instruments of the Czech Academy of Sciences. The high-power Verdi-V6 (Coherent, USA) laser provided up to 6 W in continuous mode at a wavelength of 532 nm and was focused into the sample through an Olympus PLN 20x objective (NA 0.4). The scattered light was collected through the same objective and separated by an Acton SP-2556 spectrometer (Teledyne Princeton Instruments, USA) featuring a 1200 gr/mm diffraction grating onto an Andor Newton DU920P Bx-DD CCD camera (Oxford Instruments Andor, UK) (**Figure S12a**). The microfluidic setup was kept as close as possible to the fluorescence configuration, except that the detection zone was formed by a glass capillary (ID 0.5 mm, OD 0.7 mm) coupled to the PFA tubing through a sleeve-and-union interface that provided a uniform internal diameter and prevented the droplets from tearing (**Figure S12b, c**). The total flow rate in the system was 600 µl/h and, of the 40 min between droplet generation and detection, 30 min were spent in a heated bath set to 37 °C. Spectra (acquisition time 0.1 s, 2 accumulations) were collected automatically whenever the water peak intensity reached a selected threshold, yielding approximately 750 spectra per enzyme ratio after discarding the first 2 minutes required for the system to settle. The laser output power was set to 6 W, corresponding to a measuring power of approximately 4.5 W at the sample. Data processing comprised smoothing by the Savitzky-Golay algorithm, cropping of the spectra to 2700-3100 cm^-1^, min-max normalization, grouping and averaging by enzyme ratio, subtraction of a processed averaged blank and polynomial background removal, followed by a non-negative least squares (NNLS)^56^ decomposition establishing the contributions of TCP, DCP and GLY. All acquisition and processing parameters are described in the **Raman detection mode** section of **Supplementary Information,** and the custom code is listed under **Code availability**.

#### Analysis of BO datasets by agentic AI systems

The analysis of BO datasets was performed by Theorema Lab 1.0.9^32^, Claude Opus 5,^35^ Deepseek V4 Flash,^36^ GLM 5.2^37^ and Kimi K3^38^. All sessions were performed between August 25-28, 2026. The input datasets, the full chat history, codes, output files and usage reports are available on Zenodo (**Data availability**), while the prompts and descriptions of the analytical methods applied by the agents are summarized in **Supplementary Analysis of CascadeMAP Experiments.** Agreement between the peak means from the automated detection script and from post-processing by each agentic system (prompt 01) was assessed manually by weighted Deming regression, with |slope − 1| ≤ 0.05 as the criterion for a satisfactory result. The regression procedure and the accounting of tokens and prices in the **Analysis of BO datasets by agentic AI systems** section of **Supplementary Information**.

## Code availability

Custom analysis code developed by the authors for this study is openly available: the RAMSPECT Raman spectral processing pipeline at https://github.com/pavlinasik/RAMSPECT, and the MAPit Bayesian-optimization module at https://github.com/loschmidt/CascadeMAP-Bayesian.

Theorema Lab 1.0.9 is a proprietary multi-agent AI system developed by Theorema and is not released as open source. Theorema Lab is currently in private beta. Access for academic peer review and methodological verification can be requested at https://theorema.ai/lab-access. Editors and referees of this manuscript were granted access on request during peer review.

## Data availability

The datasets collected in this study were deposited in Zenodo under the following entries. The raw and processed data from robustness test of CascadeMAP (**Figure S5**) are available on doi: 10.5281/zenodo.22898582, <u>preview link</u>. The raw and processed data from DoE experiments on the glycerol detection pathway (**Figure 2c,e**) are available at doi: 10.5281/zenodo.19001049, <u>preview link</u>. The raw and processed data from the DoE experiment on TCP degradation pathway using fluorescence detection (**Figure 3c**) are available at doi: 10.5281/zenodo.22931847, <u>preview link</u>. The raw and processed data from DoE experiment on the TCP degradation pathway using Raman detection (**Figure 3d**) are available at doi: 10.5281/zenodo.22900246, <u>preview link</u>. The raw and processed data from BO of the glycerol detection pathway in terms of temperature, pH and enzyme ratios (**Figure 4c**) are available at doi: 10.5281/zenodo.22931751, <u>preview link</u>. The input and output data for agentic AI systems (**Figure 4e,f** and **Figure 5**) are available at doi: 10.5281/zenodo.22816000, <u>preview link</u>.

## Acknowledgments

The authors would like to acknowledge funding from the Czech Ministry of Education (grants CLARA CZ.02.01.01/00/23_029/0008437, INBIO CZ.02.1.01/0.0/0.0/16_026/0008451, TEAMING CZ.02.1.01/0.0/0.0/17_043/0009632, ESFRI RECETOX LM2018121, ESFRI ELIXIR LM2018131, EXCELES Oncology LX22NPO5102). The authors also acknowledge financial support from the Swiss National Science Foundation (grant no. 205321_176011) and ETH Zurich. The authors acknowledge financial support from the Czech Science Foundation (project no. 25-15784L). This project has received funding from the European Union’s Horizon 2020 research and Innovation program (TEAMING 857560 and Sinfonia 814418). The article reflects the author’s view, and the Agency is not responsible for any use that may be made of the information it contains. Michal Vasina acknowledges the financial support of his doctoral study by the scholarship Brno Ph.D. Talent. Electron microscopy and Raman spectroscopy analysis provided at Core Facility Electron microscopy and Raman spectroscopy, ISI CAS, Brno, CZE, is supported by MEYS CR (LM2023050 Czech-BioImaging).

## Competing interests

Michael Jirasek, Filip Dousek and Hynek Walner are employees of Theorema Inc., which develops and commercializes the multi-agent AI platform used in this study. The remaining authors declare no competing interests.

## Notes

### Summary of Updates

This version of the manuscript has been revised to include a systematic comparison of five agentic AI systems using two active learning datasets comprising 2.1 GB of raw data. The systems were evaluated on autonomous data processing, multivariate analysis and reporting. Three systems completed all tasks, whereas two failed at the first task. The revised manuscript discusses the capabilities and limitations of these systems and the need for human oversight before AI systems are entrusted with direct instrument control. We also added a comparison of CascadeMAP with published self-driving experimentation platforms, considering only experiments actually performed. Among the enzyme-focused platforms compared, CascadeMAP uses the smallest aqueous-phase volume per condition (approximately 12 uL), achieves the highest measured throughput (approximately 650 to 1,300 conditions per day) and completes the largest number of closed-loop optimization cycles (100). In addition, the manuscript text has been streamlined to improve readability and clarity. These revisions do not change the interpretation of the results or the conclusions of the study.

