## Supplementary information for "CascadeMAP: Autonomous Closed-loop Optimization of Enzyme Cascades via Microfluidics, Machine Learning and Agentic AI"

Claude Opus 5 ... 36
Deepseek V4 Flash ... 38
GLM 5.2 ... 40
Kimi K3 ... 40
Theorema Lab 1.0.9 ... 41
Supplementary References ... 43

#### Supplementary Figures and Tables

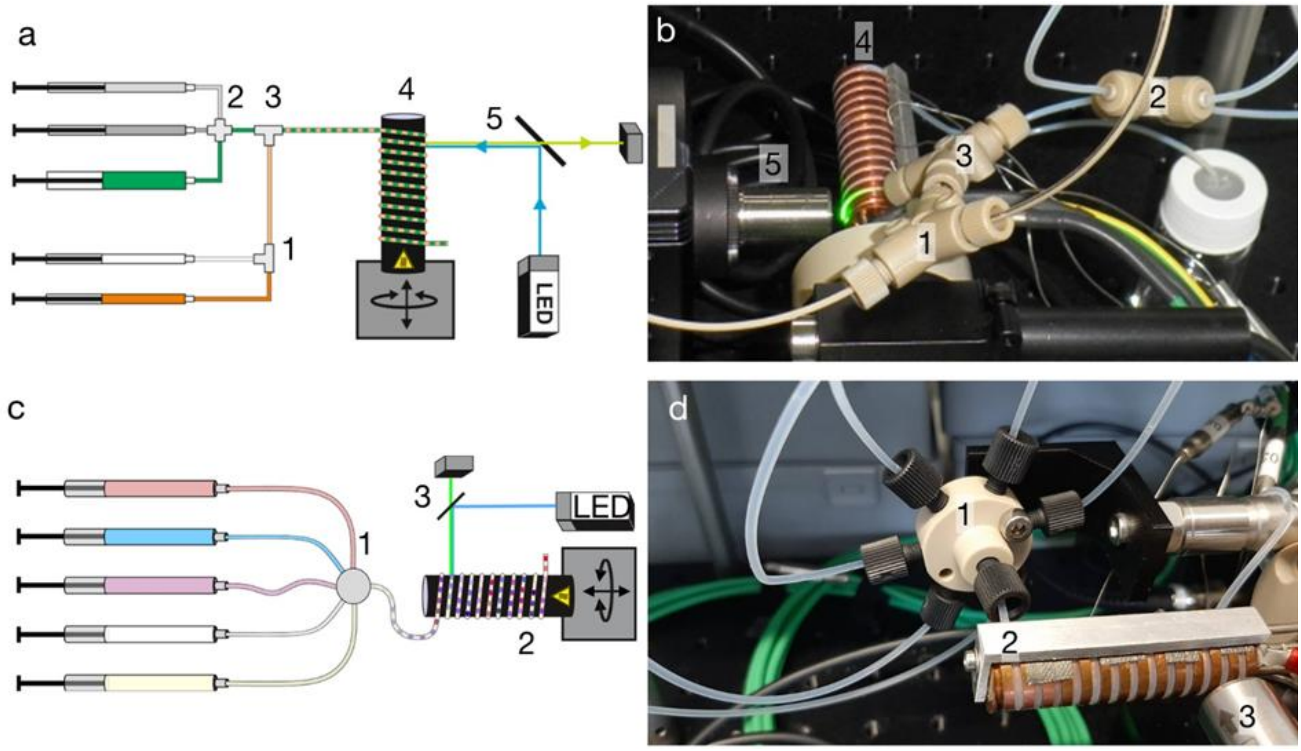

**Figure S1 | Comparison of two strategies for droplet generation within a microfluidic platform. a-b,** Scheme and photo of CascadeMAP setup for the sequential mixing of individual aqueous phases (on Y-shaped connectors - 1, and cross - 2) before droplet generation (3). After being generated, the droplets travel to a temperature-controlled rod (4) and are quantified by fluorescence measurements (5). **c-d**, Scheme and photo of the current setup for mixing all phases and droplet generation on a single manifold (1). The reaction incubator (2) and fluorescence detection (3) remain the same as in a-b.

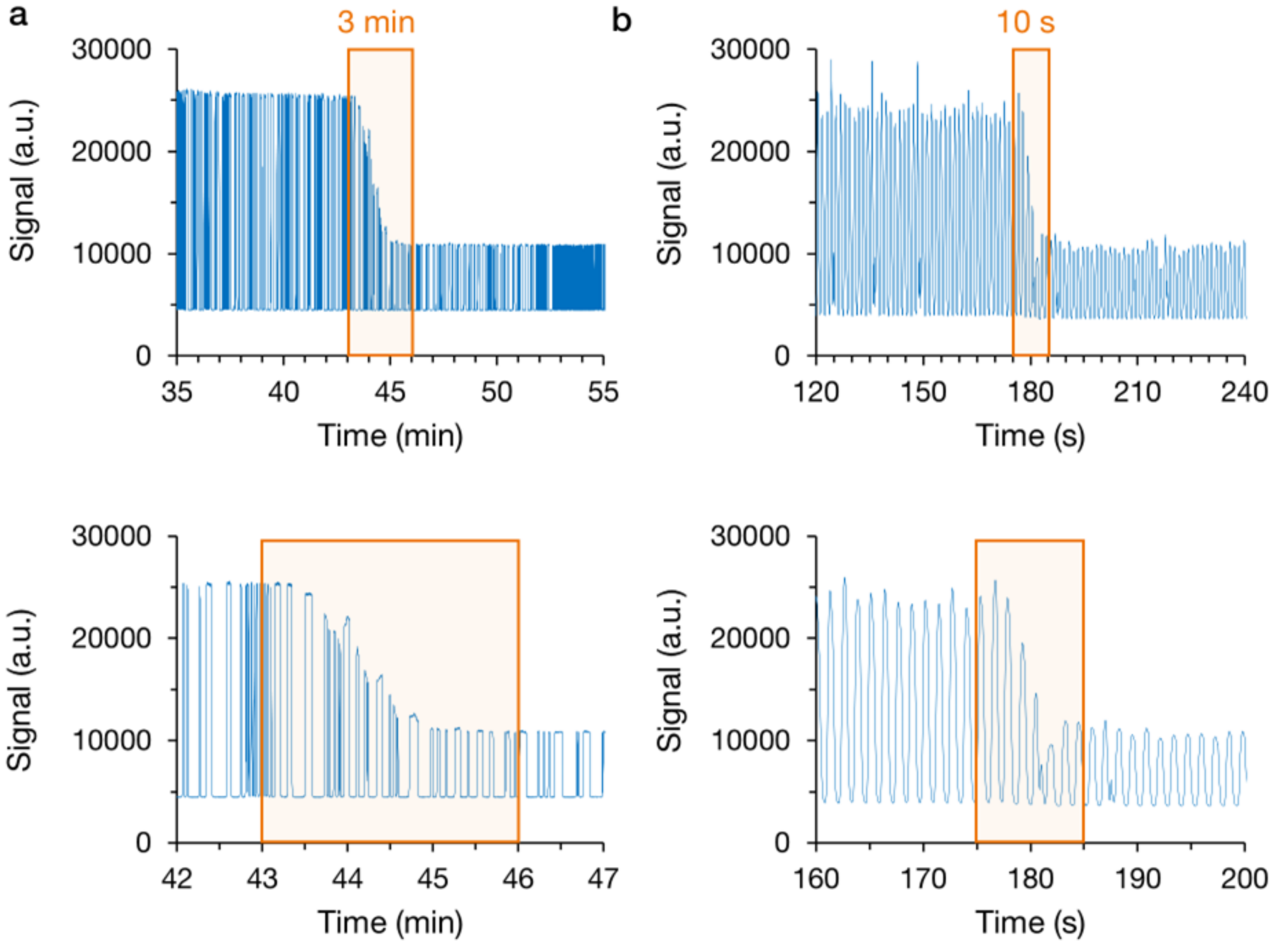

**Figure S2 | Dispersion effects when changing flowrate combinations.** **a**, Mixing of aqueous phases prior to the droplet generator, **b**, simultaneous mixing and droplet generation on the manifold. The signal in both images shows quenching of the HPTS dye when mixed with HCl. The orange square top corresponds to the same square at the bottom.

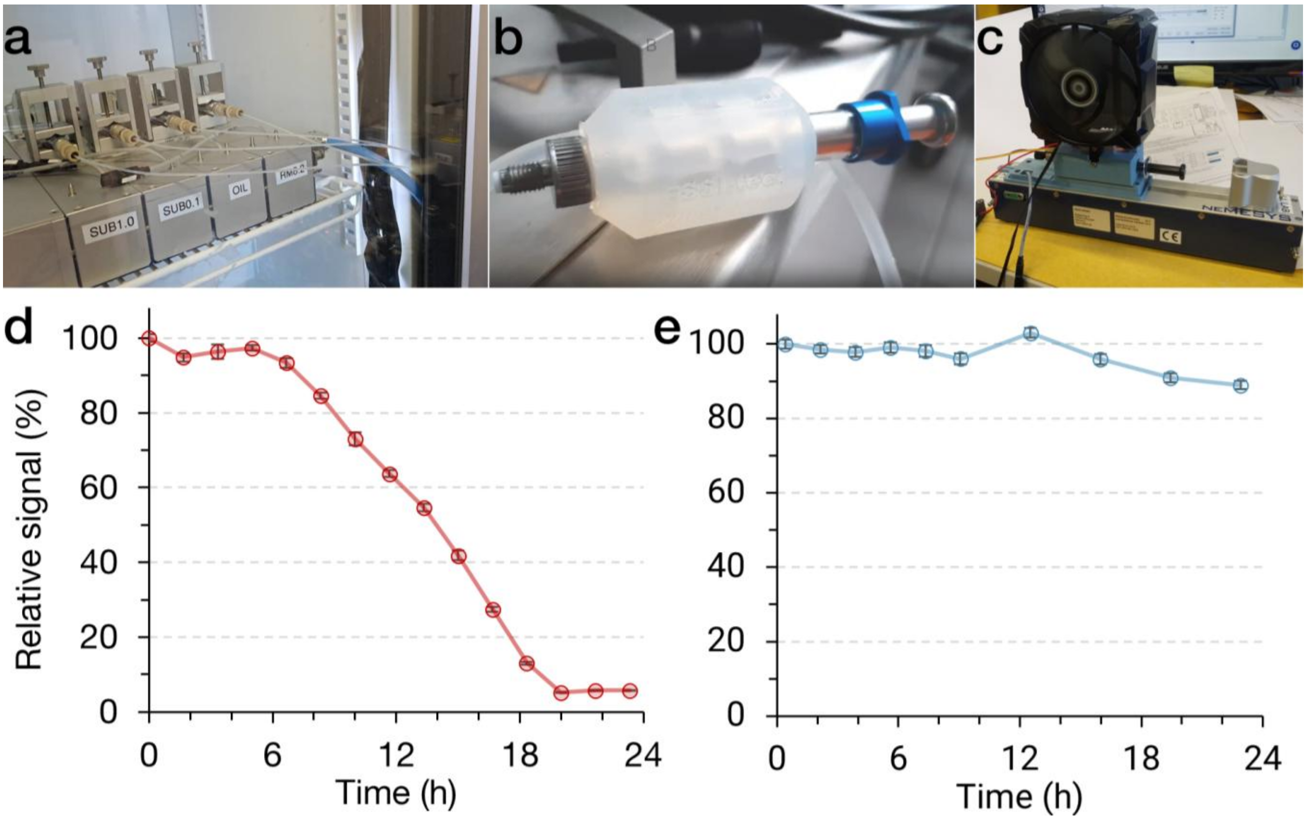

**Figure S3 | Temperature-controlled cooling within syringes. a-c**, The systems for maintaining low temperature of samples in syringes, including (**a**) a refrigerator, (**b**) a syringe sleeve (Essi Tech, Slovenia), and (**c**) a stainless-steel Nemesys syringe pump with an in-house fabricated Peltier-based temperature-controlled syringe holder **d-e**, Signal change when keeping enzyme samples (**d**) at room temperature and (**e**) in the refrigerator. Both graphs show the running of the glycerol detection pathway over about 24 hours.

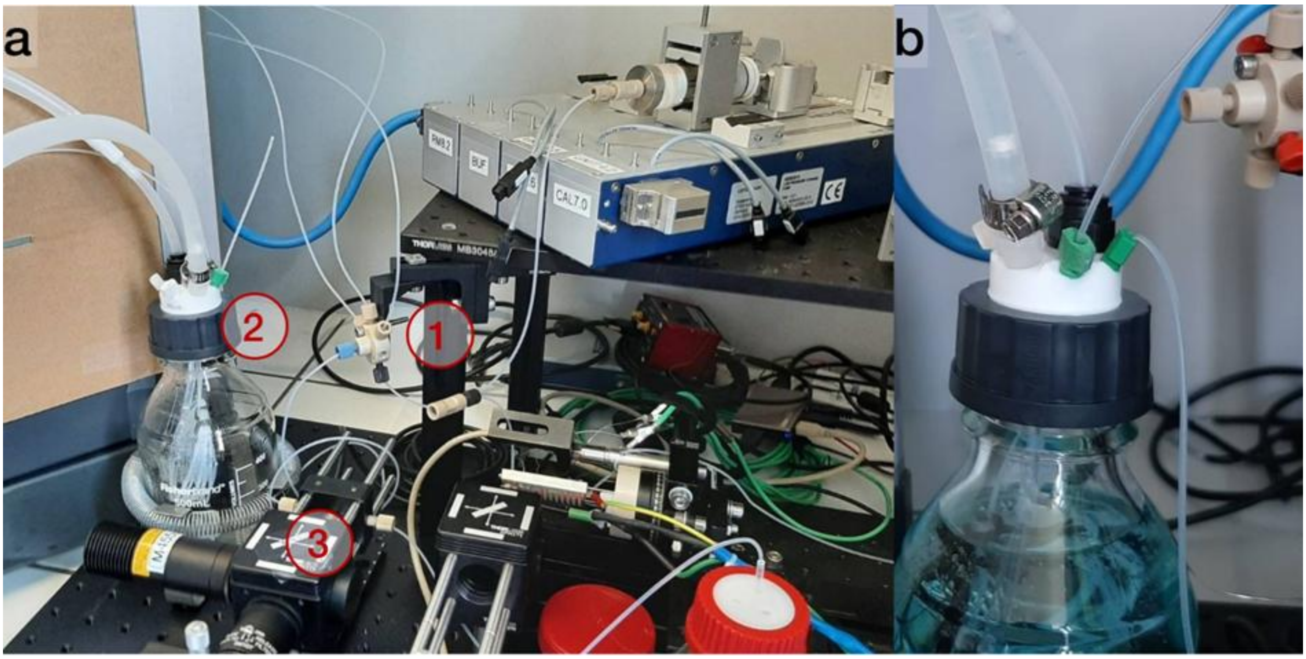

**Figure S4 | Heated reaction zone for longer reaction times. a**, CascadeMAP platform with droplet generator (1), temperature-controlled reaction zone (2), and end-point detection cell (3). **b**, Close-up of the reaction zone; the flask is filled with a temperature-regulating liquid providing both heating and cooling.

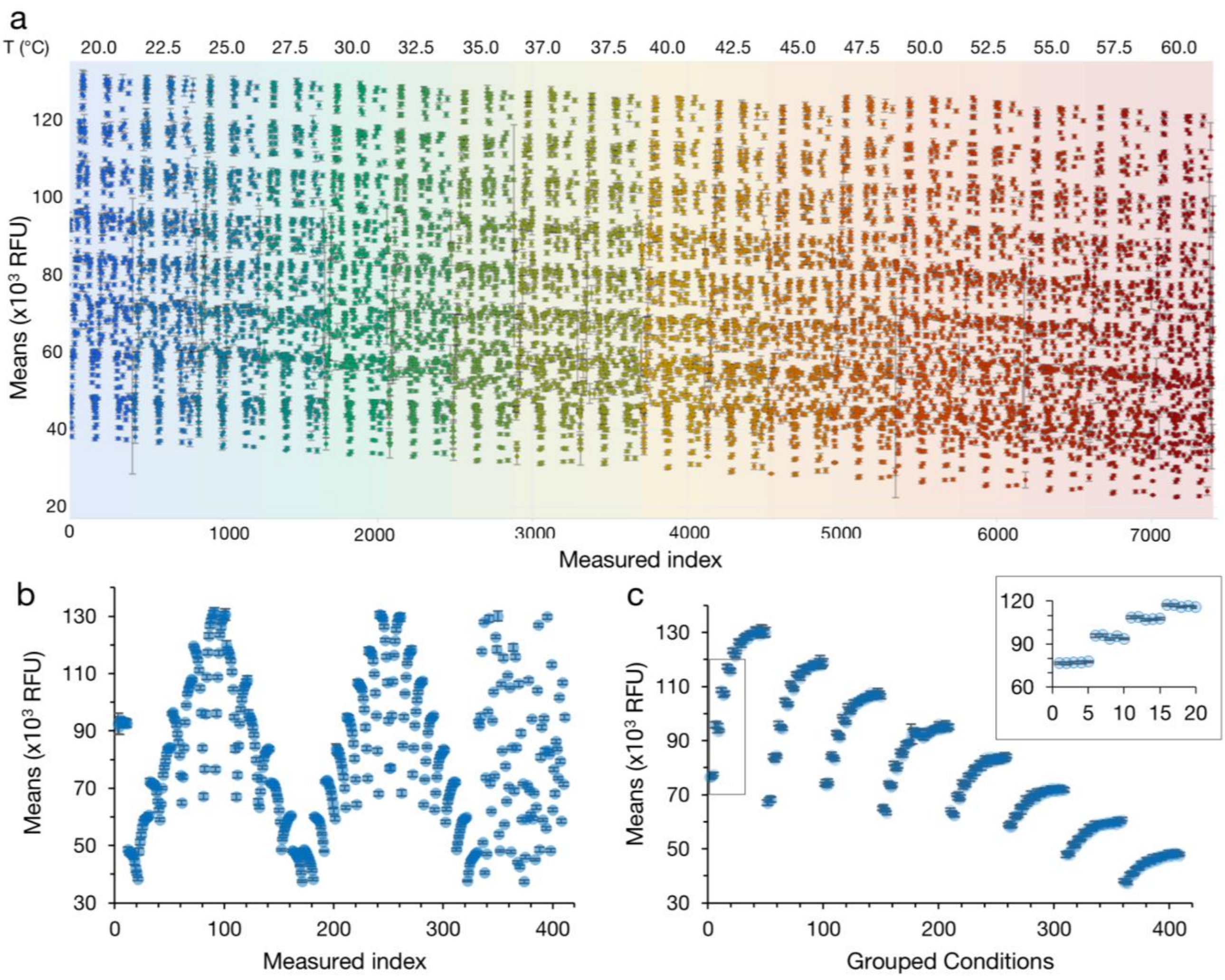

**Figure S5 | The robustness test of CascadeMAP.** The figure presents results from a DoE robustness run of about 7 days of unattended operation: 7,398 condition measurements on >295,000 droplets. The experiment tested the pH and temperature dependence of fluorescence dye HPTS and also the effect of its concentration. **a**, Means of all measured conditions are color coded based on increasing temperature from 20 to 60 °C (blue to red). The exact temperatures are displayed above the chart. **b**, Representative excerpt of data measured at 20°C. In each temperature, 411 conditions were measured in total. These include 10 equilibration conditions of equal flow rates of HPTS, Tricine and BTP, 1 auxiliary condition and 5 replicates of 80 conditions of varying all three components. Each replication of the 80 conditions is measured in a particular order: Minimized flow rate changes in between conditions going up and down with HPTS concentration for the first two replicates (measured index 11-170), partial minimization of flow rate changes with jumps from high tricine flow rate to low tricine flow rate (measured index 171-330) and finally random order of conditions (330-410). **c**, Each five replicates of a particular condition were grouped and ordered based on decreasing HPTS flow rate, then decreasing tricine and finally decreasing BTP. The inset shows in detail the first 20 replicates corresponding to 4 different conditions (marked by the rectangle with black outline).

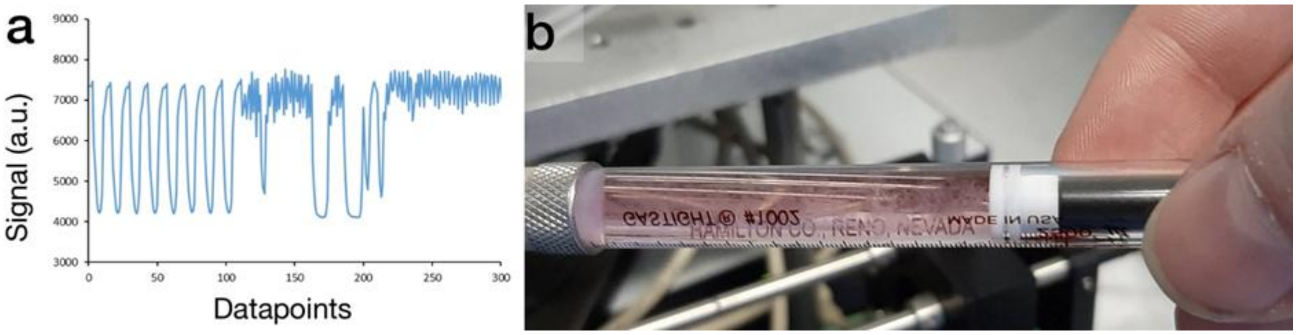

**Figure S6 | Amplex Ultra Red (AUR) behavior on CascadeMAP. a**, Impaired droplet generation due to AUR precipitation, limiting its suitability for long-duration CascadeMAP experiments. **b**, Precipitation of AUR observed ~30 minutes after syringe loading.

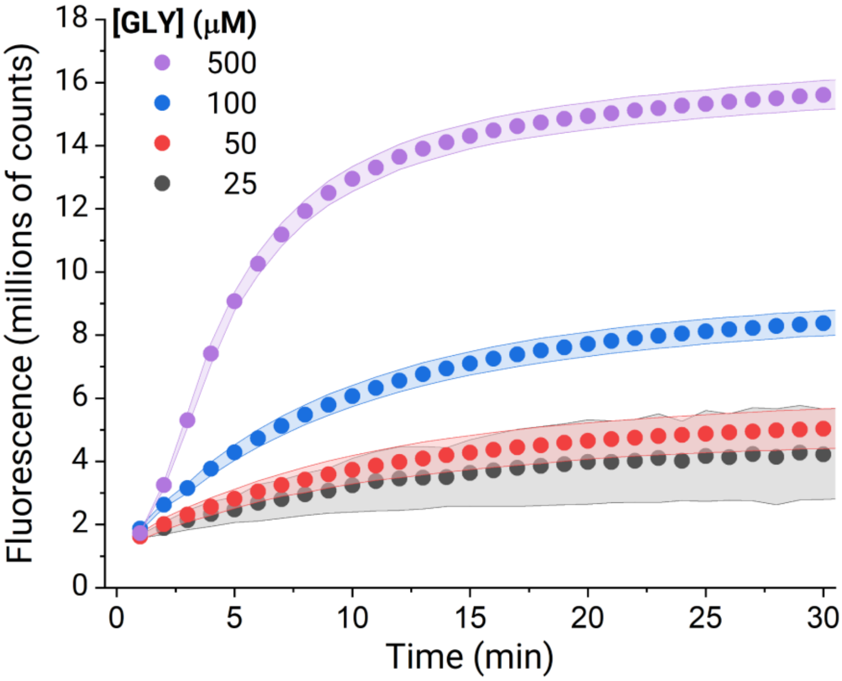

**Figure S7 | Testing of Aminophenyl fluorescein (APF) on the MTP reader.** Various concentrations of glycerol were tested using the GLY detection pathway composed of enzymes in a 1:1:1 ratio, at a total activity of 1 U/mL. The measurements were done in triplicate.

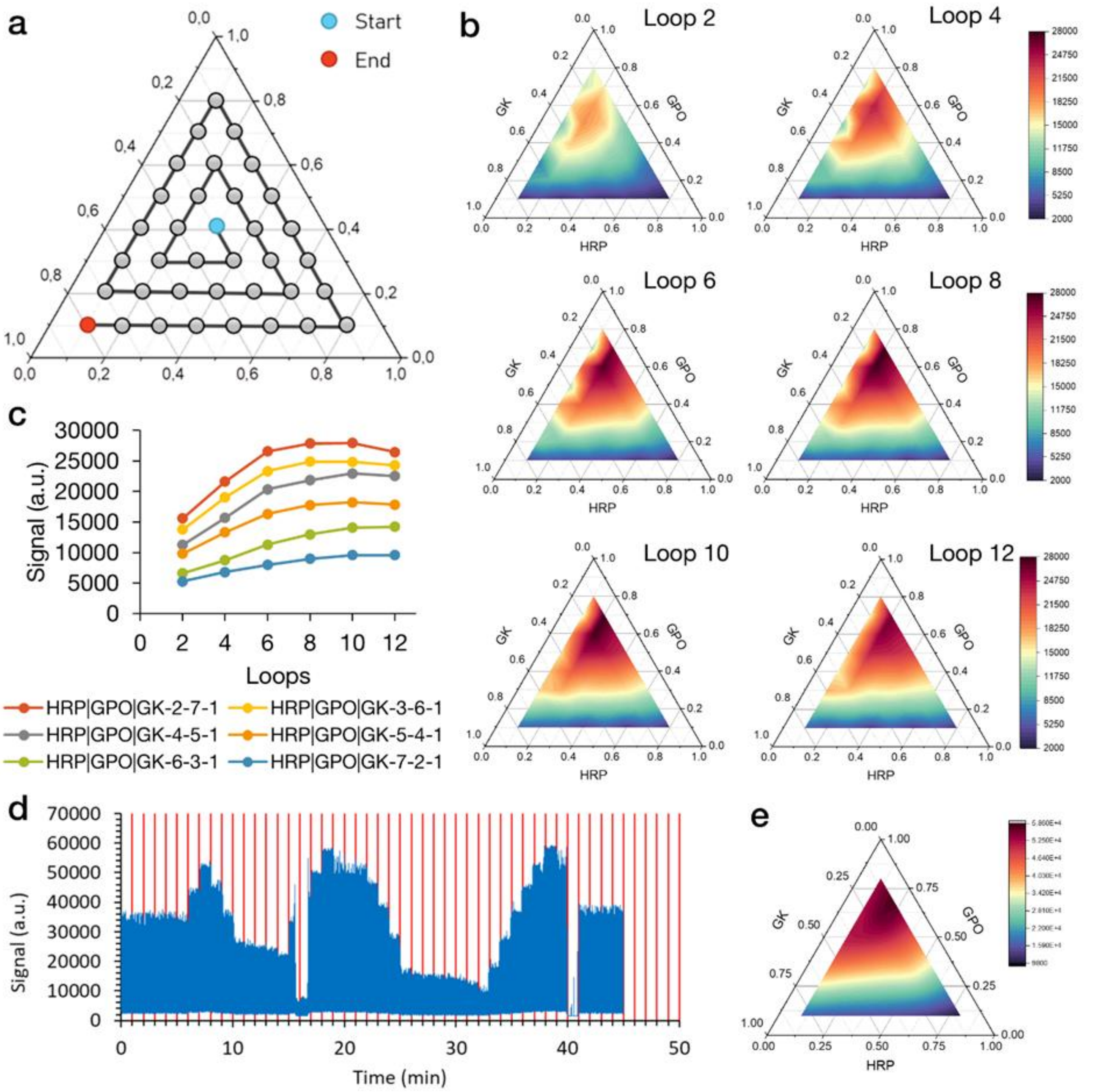

**Figure S8 | Optimization of enzyme ratios within the glycerol detection pathway. a**, The design of the experiment is based on a ternary plot diagram, where each axis represents the ratio of a single enzyme. The graph also shows the sequence of the measurement from the start (middle, blue point) to the end (bottom left, red point). **b**, The resulting ternary plots for six loops (reaction times) from the optimization experiment being run in the safe time-resolved detection mode. **c**, Time-resolved data shown for 6 different enzyme ratios as indicated in the legend. **d**, The raw data of end-point detection, where each condition is being changed every minute (red lines). The data was recorded in Loop 12. **e**, The ternary plot based on an endpoint detection experiment.

**Table S1 | Comparison of "safe time-resolved" and "stepwise end-point" data acquisition modes.** Both approaches are compared for a 3-parameter optimization of enzyme ratios, testing 36 enzyme ratios.

| Mode | Time points | Efficiency (%)[1] | Total experimental time (hours) | Total Enzymes consumption (µg) |
|---|---|---|---|---|
| Stepwise end-point | 1 | 100 | 0.66 | 7.7 |
| Safe time-resolved | 6 | 50 | 7.2 | 83 |

[1] Efficiency means the ratio of detected vs. generated droplets

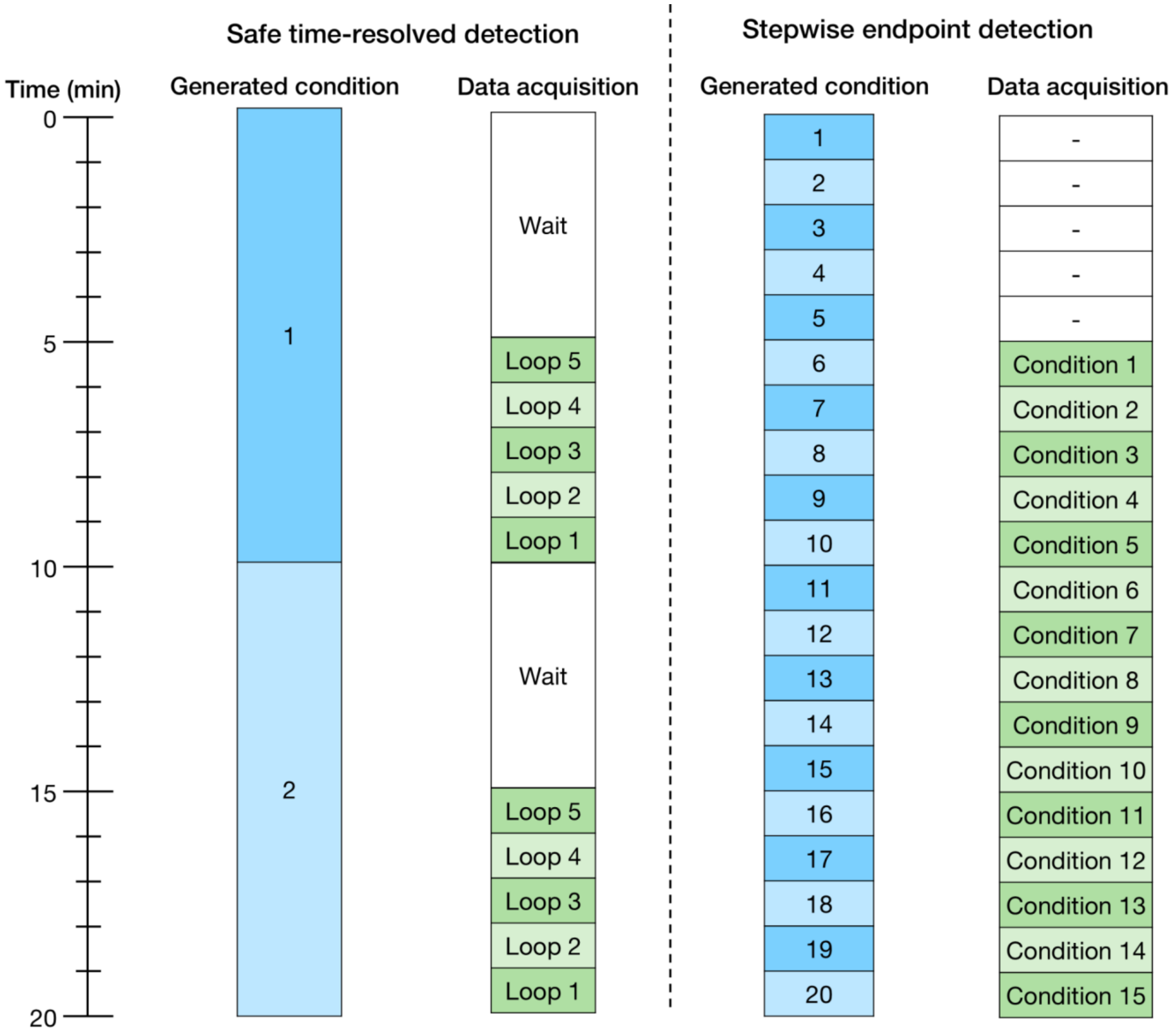

**Figure S9 | Comparison of safe time-resolved and stepwise endpoint detection modes.**The scheme shows the process of data collection for safe time-resolved (left) and stepwise endpoint (right) detection mode. On the time scale of 20 minutes with reaction time of 5 minutes, the generated conditions data acquisition is shown by blue and green shadowed tables, respectively.

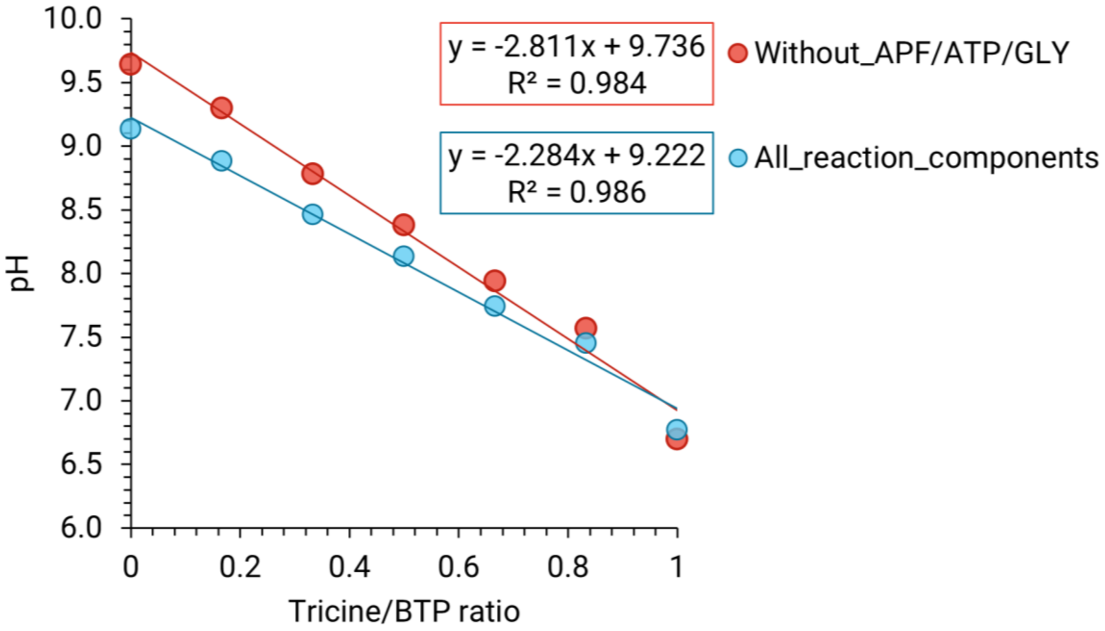

**Figure S10 | pH calibration accounting for reaction components.** The graph shows linear regressions of pH dependence on Tricine/BTP ratio of the buffer mixture (the buffer mixture forms 25% of the total reaction mixture). The red points show the mixture without added GLY/ATP and APF, while the blue points denote reaction in progress, assuming no major pH changes since the buffer concentrations are high. The data were measured at room temperature (~25°C).

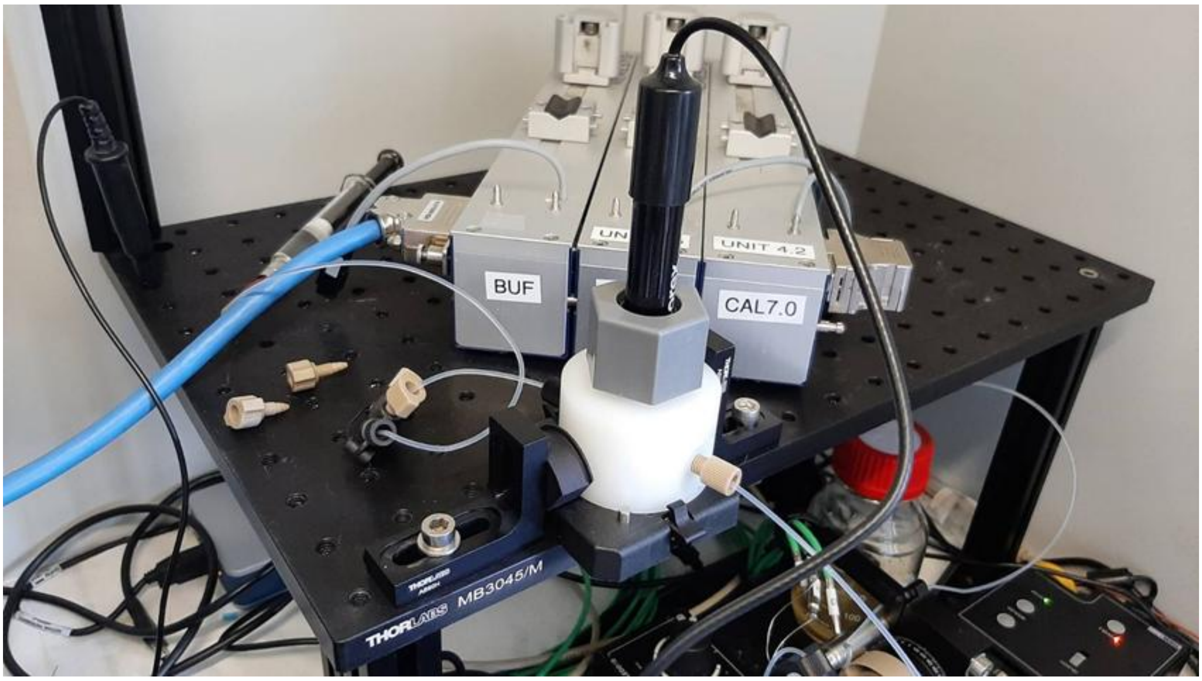

**Figure S11 | Online pH measurement.** The photo of a flow cell with a pH electrode to monitor the pH of a flowing mixture of Tricine and Bis-Tris propane.

**Table S2 | Offline pH calibration**. The pH was measured for 4 enzyme ratios (defining the ternary plot of enzyme ratios) and for temperature as indicated above the individual heat maps.

30 °C

| Tric<br>BTP | 0.00%<br>100.00% | 16.67%<br>83.33% | 33.33%<br>66.67% | 50.00%<br>50.00% | 66.67%<br>33.33% | 83.33%<br>16.67% | 100.00%<br>0.00% |
|---|---|---|---|---|---|---|---|
| HRP 80%<br>GPO 10%<br>GK 10% | 9.00 | 8.60 | 8.32 | 8.02 | 7.71 | 7.36 | 6.75 |
| HRP 10%<br>GPO 80%<br>GK 10% | 9.00 | 8.67 | 8.23 | 7.92 | 7.64 | 7.34 | 6.69 |
| HRP 10%<br>GPO 10%<br>GK 80% | 8.98 | 8.72 | 8.30 | 8.02 | 7.67 | 7.38 | 6.69 |
| HRP 33%<br>GPO 33%<br>GK 33% | 8.85 | 8.69 | 8.36 | 8.09 | 7.64 | 7.43 | 6.74 |

35 °C

| Tric<br>BTP | 0.00%<br>100.00% | 16.67%<br>83.33% | 33.33%<br>66.67% | 50.00%<br>50.00% | 66.67%<br>33.33% | 83.33%<br>16.67% | 100.00%<br>0.00% |
|---|---|---|---|---|---|---|---|
| HRP 80%<br>GPO 10%<br>GK 10% | 8.91 | 8.55 | 8.24 | 7.84 | 7.57 | 7.23 | 6.67 |
| HRP 10%<br>GPO 80%<br>GK 10% | 8.92 | 8.59 | 8.22 | 7.84 | 7.56 | 7.23 | 6.58 |
| HRP 10%<br>GPO 10%<br>GK 80% | 8.91 | 8.64 | 8.20 | 7.94 | 7.56 | 7.25 | 6.55 |
| HRP 33%<br>GPO 33%<br>GK 33% | 8.81 | 8.63 | 8.24 | 8.00 | 7.52 | 7.31 | 6.59 |

37 °C

| Tric<br>BTP | 0.00%<br>100.00% | 16.67%<br>83.33% | 33.33%<br>66.67% | 50.00%<br>50.00% | 66.67%<br>33.33% | 83.33%<br>16.67% | 100.00%<br>0.00% |
|---|---|---|---|---|---|---|---|
| HRP 80%<br>GPO 10%<br>GK 10% | 8.89 | 8.58 | 8.19 | 7.82 | 7.56 | 7.17 | 6.48 |
| HRP 10%<br>GPO 80%<br>GK 10% | 8.80 | 8.50 | 8.12 | 7.78 | 7.45 | 7.12 | 6.49 |
| HRP 10%<br>GPO 10%<br>GK 80% | 8.85 | 8.62 | 8.13 | 7.77 | 7.44 | 7.07 | 6.44 |
| HRP 33%<br>GPO 33%<br>GK 33% | 8.78 | 8.60 | 8.20 | 7.88 | 7.49 | 7.18 | 6.66 |

40 °C

| Tric<br>BTP | 0.00%<br>100.00% | 16.67%<br>83.33% | 33.33%<br>66.67% | 50.00%<br>50.00% | 66.67%<br>33.33% | 83.33%<br>16.67% | 100.00%<br>0.00% |
|---|---|---|---|---|---|---|---|
| HRP 80%<br>GPO 10%<br>GK 10% | 8.82 | 8.38 | 8.11 | 7.71 | 7.44 | 7.06 | 6.53 |
| HRP 10%<br>GPO 80%<br>GK 10% | 8.83 | 8.46 | 8.07 | 7.66 | 7.41 | 7.06 | 6.40 |
| HRP 10%<br>GPO 10%<br>GK 80% | 8.77 | 8.51 | 8.04 | 7.75 | 7.41 | 7.08 | 6.40 |
| HRP 33%<br>GPO 33%<br>GK 33% | 8.65 | 8.51 | 8.04 | 7.84 | 7.41 | 7.14 | 6.44 |

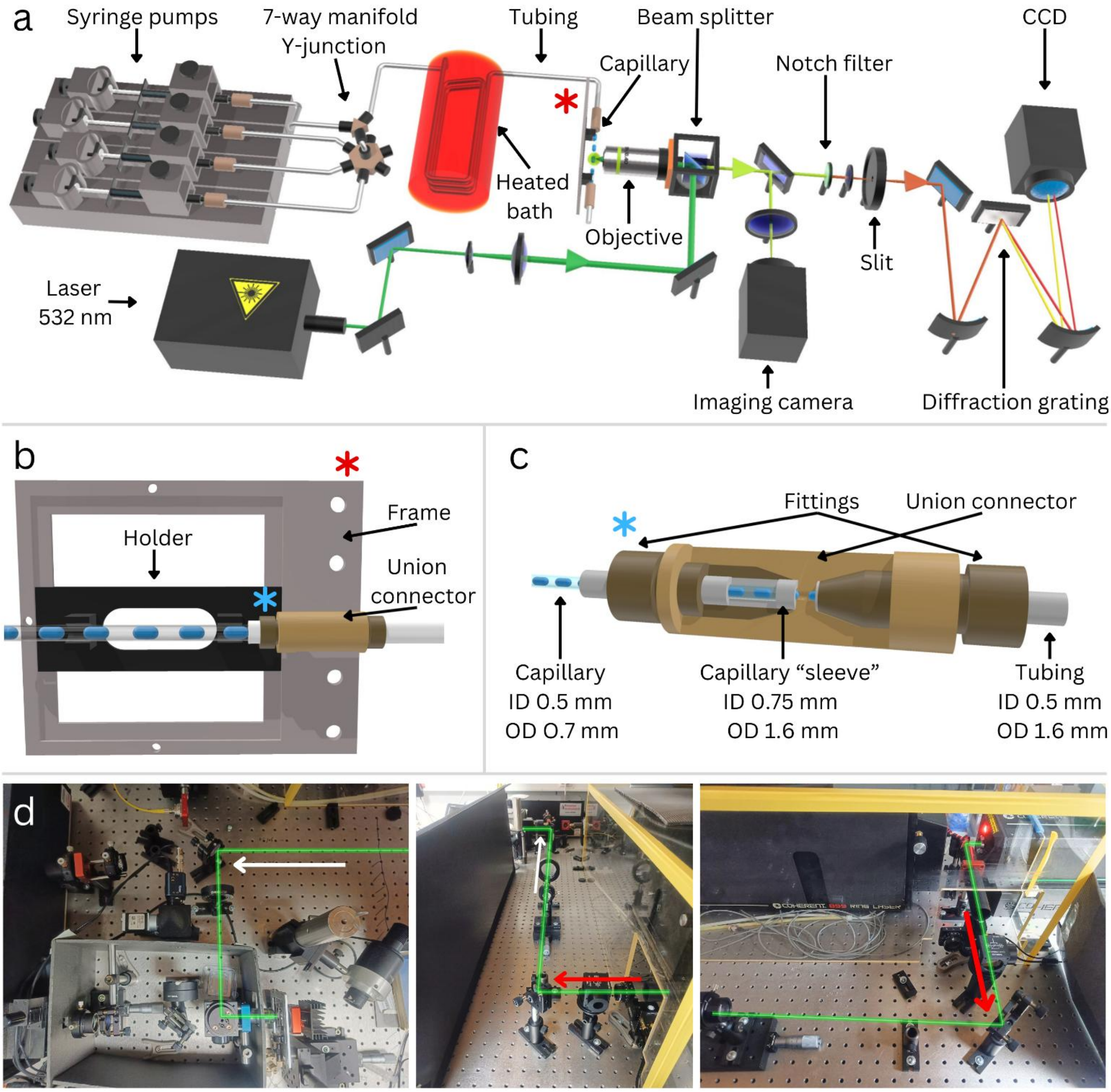

**Figure S12 | Analytical setup for the Raman detection of TCP pathway metabolites. a**, The scheme of the microfluidic platform with the Raman microspectrometer. Metabolites can be detected in the capillary. The red asterisk marks the frame supporting the holder with a capillary. **b**, Close-up of the milled frame and the 3D-printed capillary holder. The blue asterisk marks the connector for tubing/capillary transition. **c**, Detailed look inside the connector and the tubing/capillary transition. **d**, Photos of the optical pathway of the in-house built Raman instrument. The green line indicates the laser, and the arrows indicate the direction of the laser. Arrows are color-coded in pairs to provide better clarity of the components' orientation.

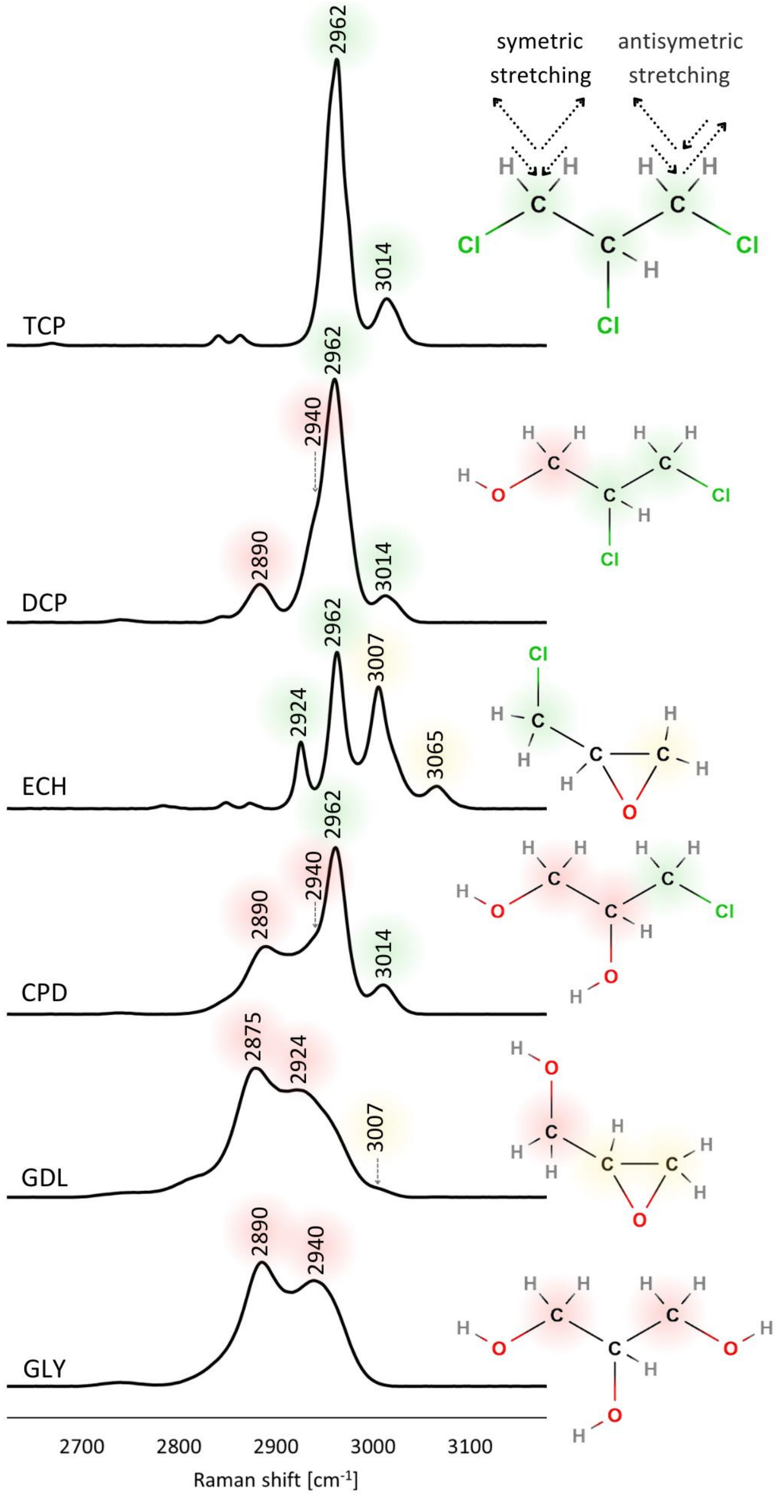

**Figure S13 | Spectra of neat compounds present in the TCP degradation pathway.** Spectra collected with InVia Renishaw for reference and calibration. Main peaks highlighted and color-matched with the respective vibration areas within the molecules. The abbreviated compound names denote: TCP - 1,2,3-trichloropropane, DCP - (*R,S*)-2,3-dichloropropane-1-ol, ECH – (*R,S*)-epichlorohydrin, CPD – (*R,S*)-3-chloropropane-1,2-diol, GDL – (*R,S*)-glycidol and GLY – glycerol.

**Table S3 | Peak assignment for all metabolites from TCP degradation.**[1–4] ν: stretching, s: symmetric, as: antisymmetric.

| Peak position [cm⁻¹] | Assignment | Metabolite |
|---|---|---|
| 2875 | $\nu CH_2(OH)_s$ | *GDL* |
| 2890 | $\nu CH_2(OH)_s$ | *DCP, CPD, GLY* |
| 2924 | $\nu CH_2(Cl)_s$ | *ECH* |
| | $\nu CH_2(OH)_{as}$ | *GDL* |
| 2940 | $\nu CH_2(OH)_{as}$ | *DCP, CPD, GLY* |
| 2962 | $\nu CH_2(Cl)_s$ | *TCP, DCP, CPD* |
| | $\nu CH(Cl)$ | *TCP, DCP* |
| | $\nu CH_2(Cl)_{as}$ | *ECH* |
| 3007 | $\nu CH_{2s}$ | *GDL, ECH* |
| 3065 | $\nu CH_{2as}$ | *ECH* |

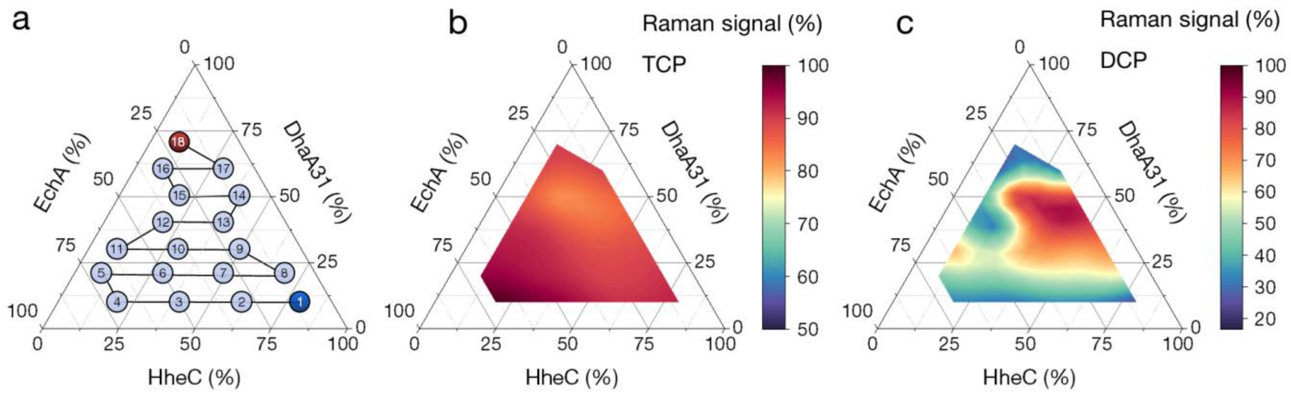

**Figure S14 | Raman-based monitoring of substrate and intermediates within the TCP pathway. a,** The design of the optimization of TCP pathway enzymes ratios using high-power Raman spectroscopy. The numbered points indicate the sequence of enzyme mixing. **b, c,** The ternary plots show relative Raman signals depending on enzyme ratios for the substrate TCP (**b**) and first intermediate DCP (**c**). The color bar for the heatmaps is shown to the right of each ternary plot. The experiment was carried out at ~30 min of reaction time, with a mean reaction temperature of ~37°C.

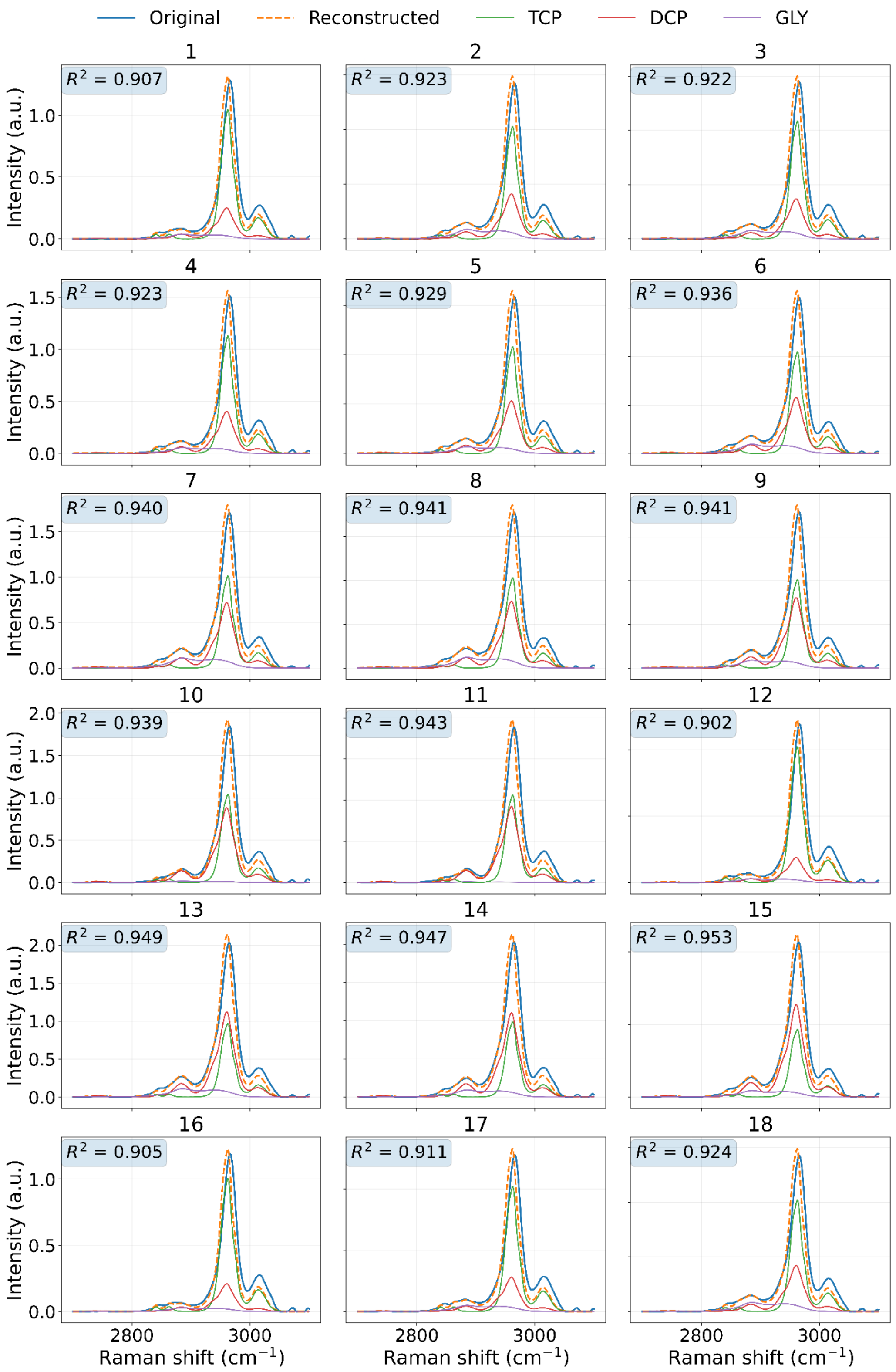

**Figure S15 | Reconstructed spectra compared to the resulting reaction spectra, including TCP, DCP, and GLY components for each enzyme ratio reaction.** Individual subplots depict a specific reaction (specific enzyme ratio) spectrum and the corresponding reconstructed spectrum created by adding TCP, DCP, and GLY component spectra calculated by NNLS. The $R^2$ value in each plot is the coefficient of determination and indicates how well the reconstructed spectra match the ground truth spectra ($R^2$ = 1.000 for perfect fits).

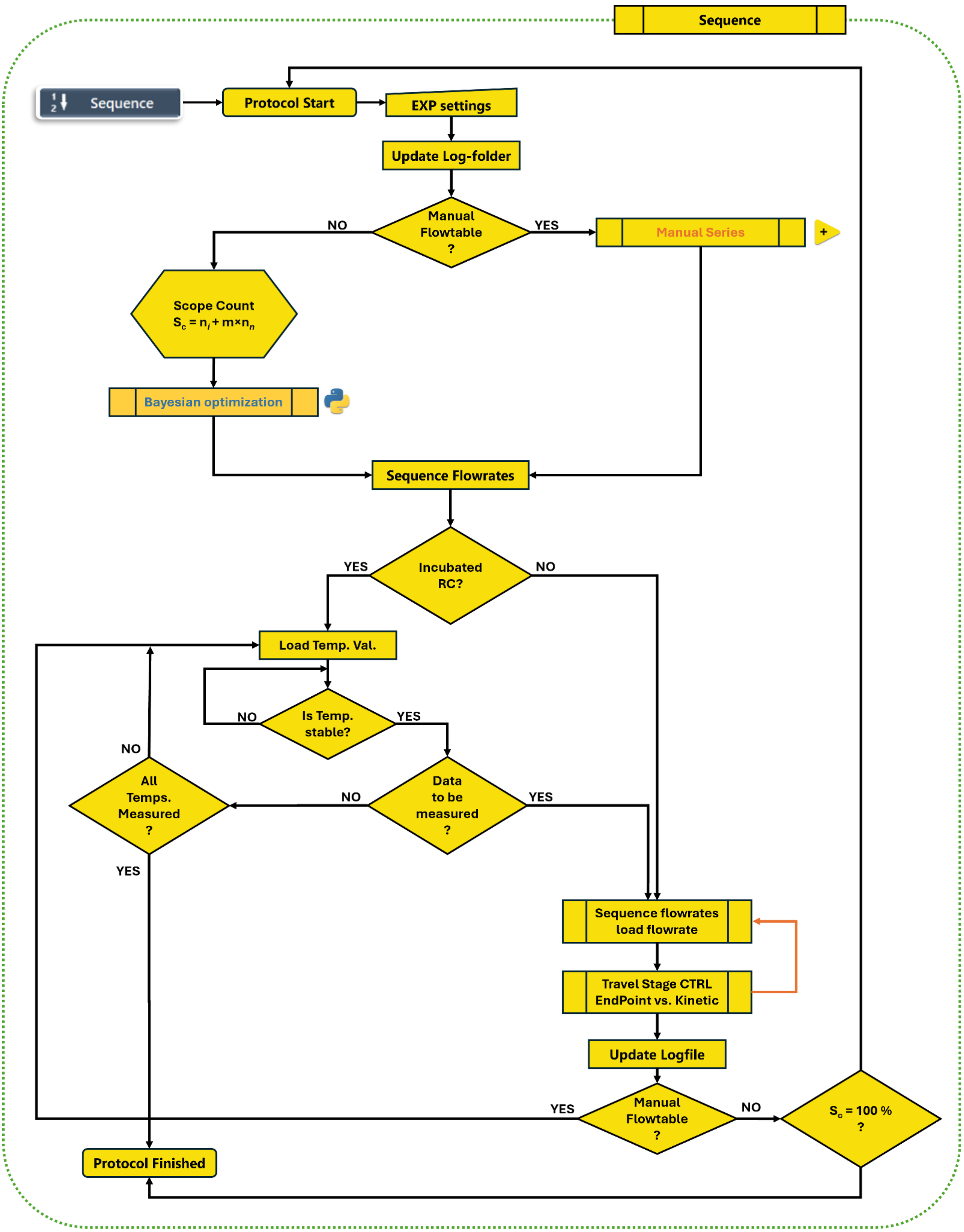

**Figure S16 | Flowchart of the sequence protocol workflow in the MAPit software.** After the user selects the protocol parameters and settings, the sequence executes automatically until all configured conditions are measured. Scope count ($S_c$) is a sum of initial points ($n_i$) and multiple (m) of predicted next points ($n_n$). The Bayesian optimization protocol runs until all scopes are measured.

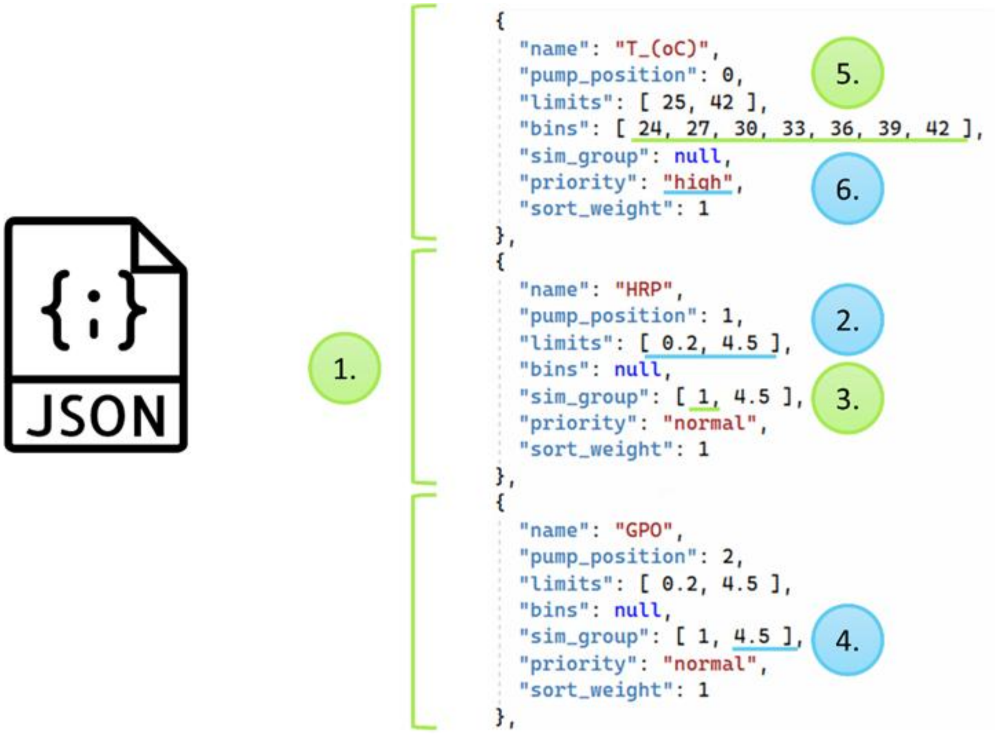

**Figure S17 | The JSON file is used to set up the BO experiment.** The beginning of the JSON file used for glycerol detection pathway optimization is shown with numbers corresponding to the list of required inputs: 1. Parameters to be shuffled ("name"; a string) correspond to a single syringe operated by a single/pump or temperature. 2. Position of the pump ("pump_position"; natural number) determines the physical position of the pump, which will be affected by shuffled parameters. 3. The values range ("limits":[min, max]; real values) within which the parameter values will be generated. 4. Discrete values ("bins"; an ascending list of real numbers) according to which the parameter values will be grouped together. 5. Groups of parameters ("sim_group":[#group_number, total_flow]; integer for the former and real value for the latter) whose sum needs to stay constant. 6. Priority ("priority"; either "high" or "normal") according to which the generated values are sorted. The value "high" is set for the temperature.

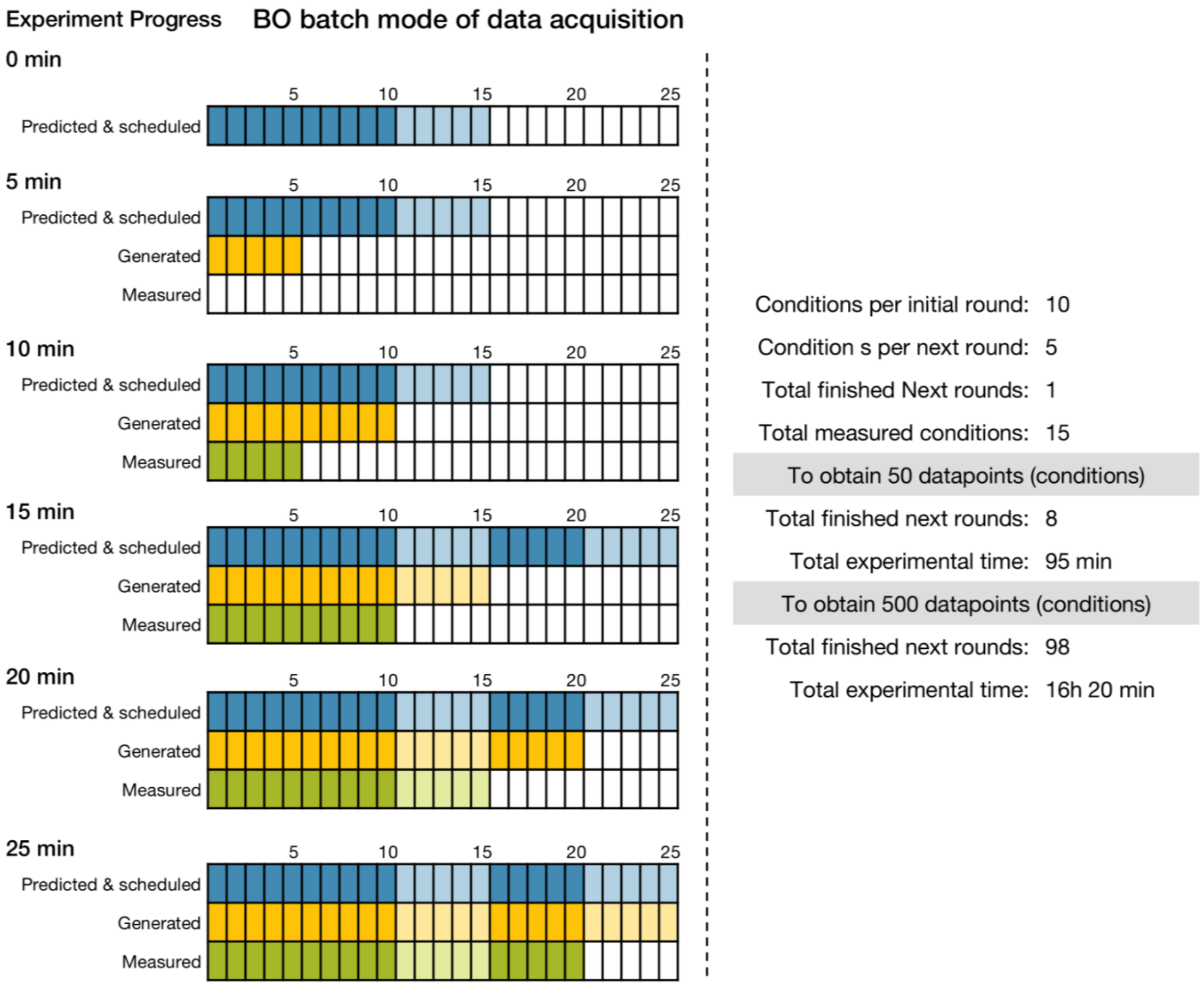

**Figure S18 | Overview of the current batch mode of data acquisition during BO experiments.** Left: The progress of an imagined experiment operated under either current batch mode and recording of predicted and scheduled conditions (blue), conditions for which droplets were generated (yellow) and measured conditions (green). Right: The statistics of the imagined experiment taking 25 minutes are supplemented by two additional extensions to obtain 50 and 500 datapoints.

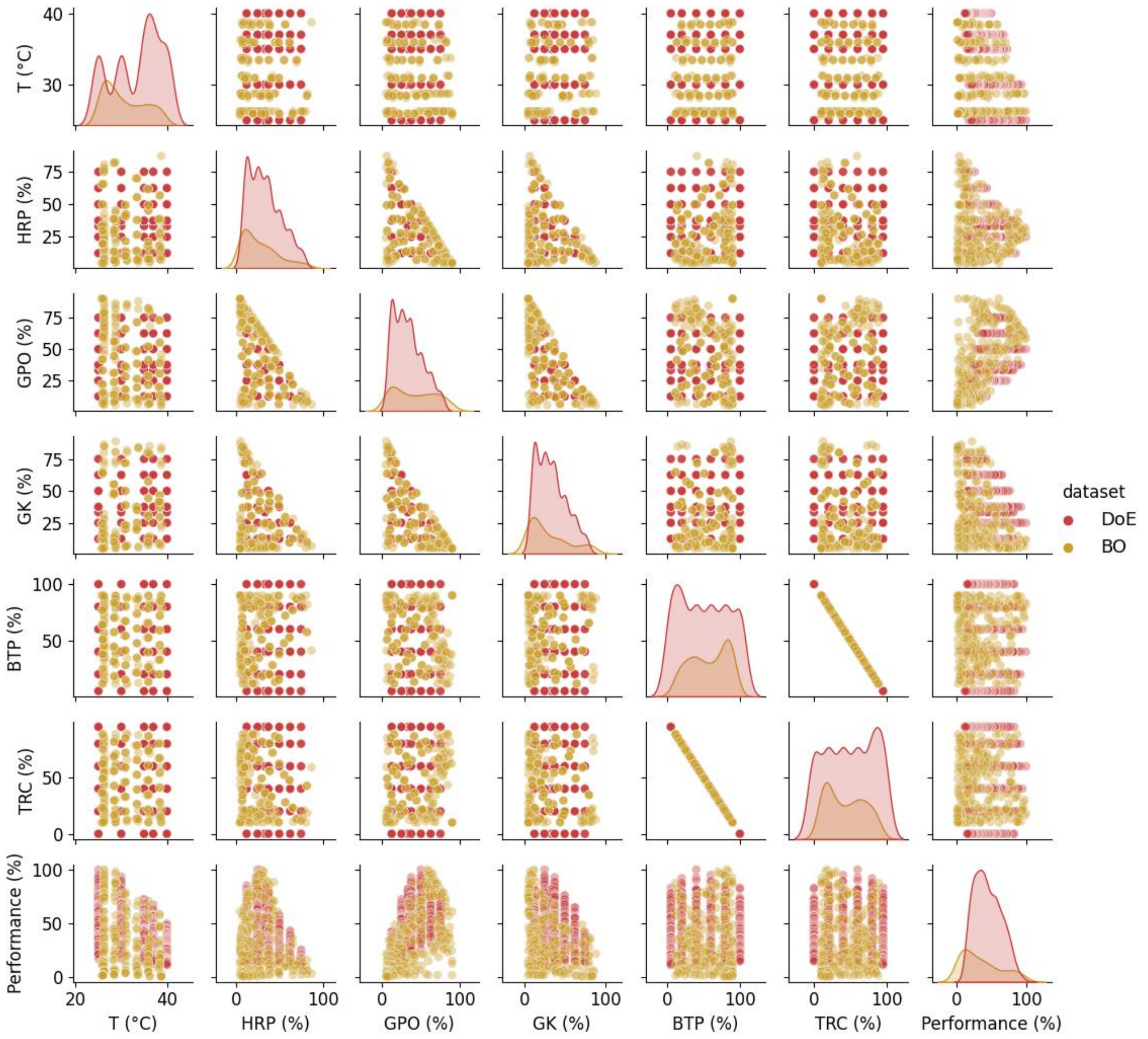

**Figure S19 | Pairplot graph for DoE and AL optimization of glycerol detection pathway.** The pairwise plot compares the data patterns for DoE and AL-based optimization of enzyme ratios, temperature and pH of the glycerol detection cascade.

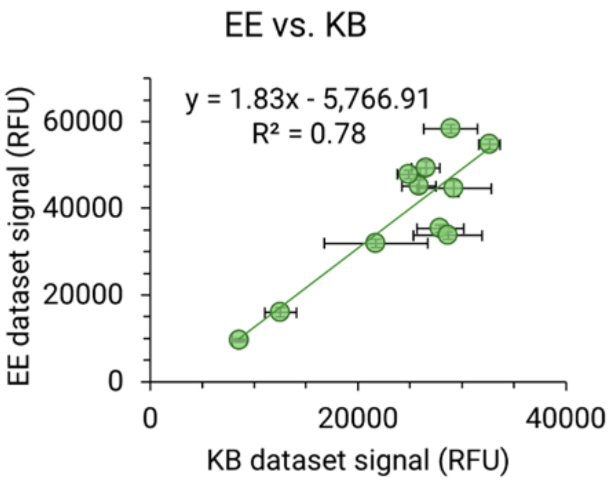

**Figure S20 | Linear regressions to compare initial round conditions for EE and KB algorithm.** Fluorescence signals and their respective SDs are plotted for EE across the KB dataset. The equation and accuracy of linear regression are shown in the plot. EE and KB are single runs; the KB data were projected onto the EE data with this regression (**Figure 4e-f**).

**Table S4 | Metrics of agentic systems' performance on autonomous data processing and interpretation of the two representative BO datasets.** Each metric (column) has 3 subcolumns distinguished by color code, where orange (first from left in each triad) denotes Claude Opus 5, light purple denotes Theorema Lab (middle in each triad) and light blue denotes Deepseek v4 Flash (first from right in each triad).

| Prompt | Wall time (min) | | | Steps (tool calls) | | | Total tokens (M) | | | Estimated price ($) | | | Necessary step-ins | | |
|---|---|---|---|---|---|---|---|---|---|---|---|---|---|---|---|
| 1 | 24 | 50 | 62 | 53 | 14 | 93 | 11.12 | 8.48 | 4.86 | 11.69 | 7.76 | 0.12 | 0 | 1 | 1 |
| 2 | 13 | 8 | 12 | 33 | 7 | 23 | 15.22 | 1.61 | 2.58 | 13.99 | 1.22 | 0.11 | 0 | 0 | 0 |
| 3 | 48 | 7 | 23 | 61 | 6 | 26 | 40.31 | 0.61 | 3.72 | 29.17 | 0.46 | 0.31 | 0 | 0 | 0 |
| 4 | 18 | 11 | 13 | 38 | 21 | 18 | 35.15 | 3.82 | 3.07 | 27.84 | 2.89 | 0.14 | 0 | 0 | 4 |
| 5 | 50 | 357 | 61 | 80 | 61 | 61 | 103.26 | 12.35 | 16.70 | 66.1 | 11.16 | 1.28 | 0 | 1 | 5 |
| **Sum** | **153** | **433** | **171** | **265** | **109** | **221** | **205.05** | **26.86** | **30.93** | **148.79** | **23.49** | **1.96** | **0** | **2** | **10** |

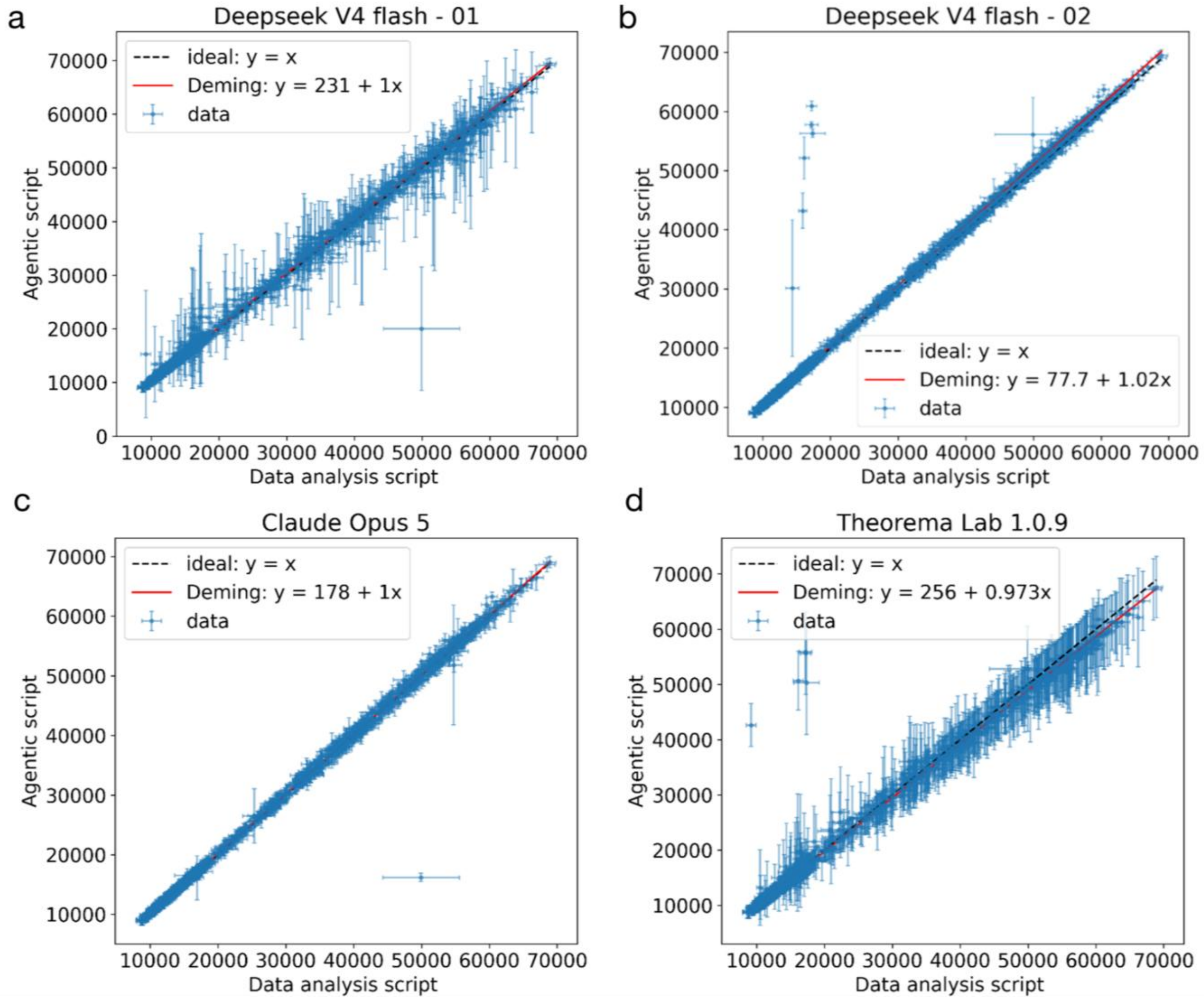

**Figure S21 | Weighted Deming regression analysis for systems that passed prompt 1. a-d,** The graphs show the agreement between the processed peaks (blue data points with corresponding standard deviations) by automated data analysis script (x-axes) and the particular agentic script (y-axes). The black dashed-line corresponds to an ideal match of both datasets, while the red solid line shows the weighted Deming regression. The Deming equation is displayed in the legend for every graph. The results of Deepseek V4 flash model show the iteration 1 (**a**) and 2 (**b**).

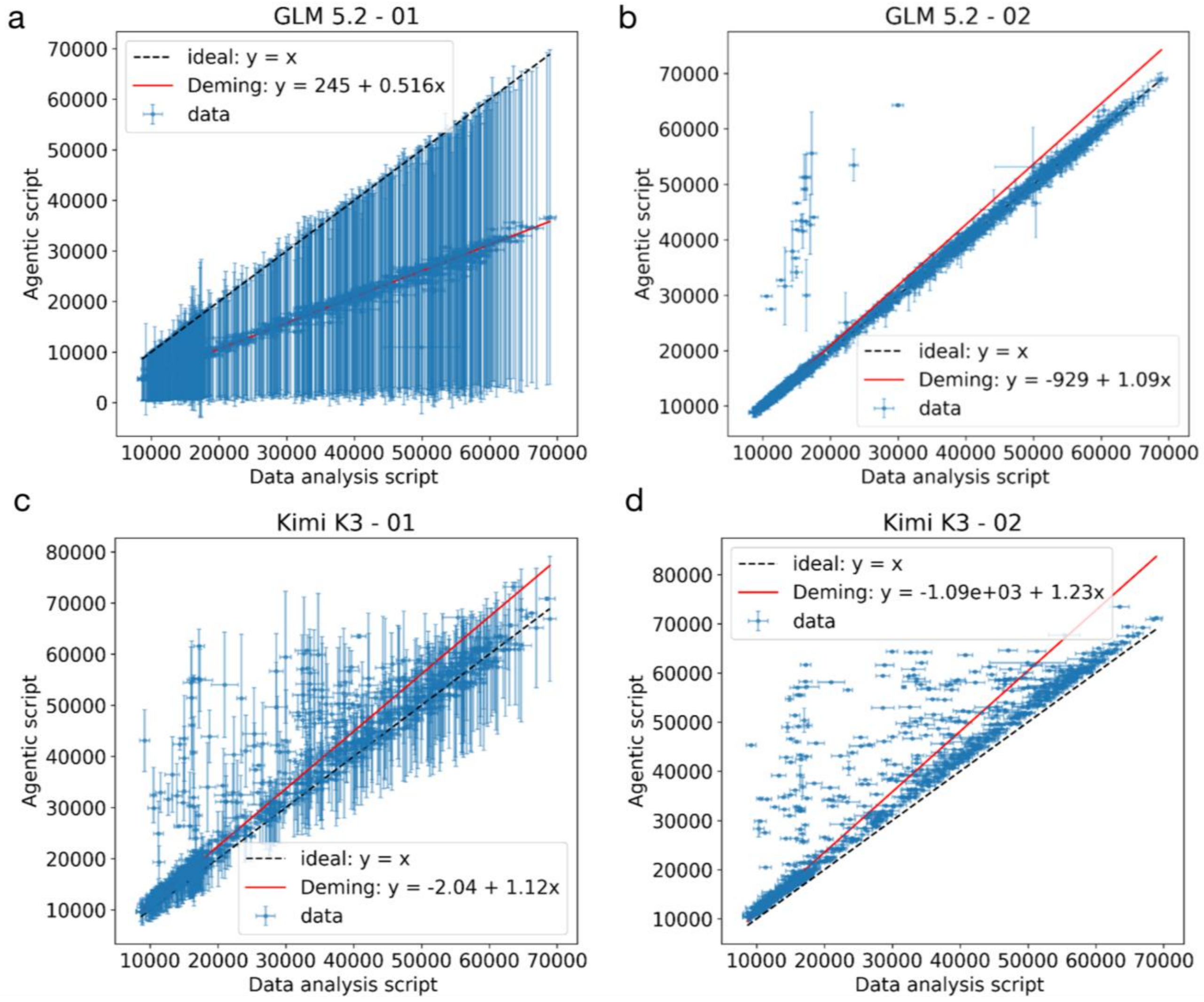

**Figure S22 | Weighted Deming regression analysis for systems that failed prompt 1. a-d,** The graphs show the agreement between the processed peaks (blue data points with corresponding standard deviations) by automated data analysis script (x-axes) and the particular agentic script (y-axes). The black dashed-line corresponds to an ideal match of both datasets, while the red solid line shows the weighted Deming regression. The Deming equation is displayed in the legend for every graph. The results of GLM 5.2 model show the iteration 1 (**a**) and 2 (**b**), while Kimi K3 also show iterations 1(**c**) and 2 (**d**).

#### Supplementary Methods

This section provides the full experimental detail underlying the condensed **Online Methods** of the main text.

##### CascadeMAP experiments with fluorescence detection

###### Hardware

The CascadeMAP hardware is based on the previously introduced and thoroughly described KinMAP.[5] Briefly, the oil and aqueous phases are loaded to GASTIGHT syringes (Hamilton, USA) and then operated by the low-pressure neMesys microfluidic pumps (Cetoni, Germany) in the positive pressure mode. All liquids travel through high-purity perfluoralkoxy (PFA) tubing 1622L (OD 1/16 inch, ID 0.02 inch, IDEX Health & Science, USA) and the droplets are generated either on 6-inlet manifold P-170 (IDEX Health & Science, USA), or on a 10-inlet manifold Z10M1PK (Vici Valco, France). For the long-term experiment, the aqueous solutions were refilled during the experiment automatically by using 3/2-way valves (Cetoni, Germany) and the oil was delivered via two 60 ml-Omnifix (Braun, Germany) syringes both mounted on a single Fusion 200 pump (Chemyx, USA). The flow of oil from these two syringes was then unified via the Y-junction P-512 (IDEX Health & Science, USA). The ratios of solutions and velocity of droplet movement were controlled by changing the flow rates of the individual pumps, keeping the constant flow rates of 12 and 12 μL/min for the aqueous and oil phases, respectively.

The droplets with the reaction mixture subsequently traveled through the tubing coiled around a copper-heating rod (diameter = 1.5 cm, length 8 cm, manufactured by PB Control, Czech Republic), which allows moving along the longer axis using a travel stage (LTS150, Thorlabs, Germany) and heating from ambient temperature up to 180 °C. The copper rod is heated by a cartridge (6.5 x 40 mm, 100 W, Farnell, Switzerland) embedded inside the rod. The temperature is monitored using a thermocouple (Sensor, Thermoelement Type K - 0.5 mm, Farnell, Switzerland), inserted into the copper block close to the surface. Temperature control is realized using a PID controller (CN7800, Omega, USA), with an observed temperature variation from the set point of 0.1 °C.

Fluorescence assays were performed using the following optical components, all purchased from Thorlabs (Germany) if it is not stated otherwise. For APF assays, excitation was provided by an LED M470L3 diode (470 nm), in combination with a DMLP505R dichroic mirror and an excitation filter XVS0470 Shortpass Filter VIS 470 nm 25 mm dia. (Asahi spectra, USA). Emission was collected at 512–518 nm range, with the LED operated at 90% intensity. For AUR assays, excitation was achieved using an LED M530L3 diode (530 nm), together with a DMLP567R dichroic mirror and a 590 nm long-pass filter (Comar 590 GY25, UK). Emission was collected in the 588–594 nm range at an LED intensity of 30%.

###### Integrated mixing and droplet generation

In typical capillary-based designs reported in the literature,[5–8] aqueous reagents are first mixed in a manifold, and droplets are subsequently generated at a downstream T-junction. While widely used, this approach introduces axial dispersion during transport from the mixing region to droplet formation. This smearing of concentration gradients compromises temporal precision and necessitates long delay times to allow for transitions between distinct reaction conditions. In contrast, CascadeMAP integrates both aqueous and oil phases within a single manifold, coupling mixing directly to droplet generation. By minimizing the dead volume between mixing and encapsulation, this integrated configuration substantially reduces dispersion and enables rapid, well-defined switching between flow rate combinations (**Figure S1, S2**), allowing precise temporal control of reagent delivery and higher fidelity in sampling, which is inherently difficult to achieve in conventional setups.

#### Maintaining sample integrity during extended experiments

Various technical solutions were implemented to preserve sample integrity and maintain stable experimental conditions during extended self-driving experiments, typically lasting more than 10 hours (**Figure S3**). These include a refrigerated enclosure, customized syringe sleeves, and specialized syringe adapters (**Figure S3a-c**). Although this setup required a larger physical footprint, the refrigerator ensured the stability of enzymes resulting in a consistent signal over time (**Figure S3d-e**). To accommodate reactions spanning longer timescales and to reach control of temperatures below room temperature, a flask-based incubator and a complementary detection cell for end-point measurements were also integrated. These additions extended the versatility of CascadeMAP, allowing it to monitor both rapid and slow enzymatic processes with equal precision (**Figure S4**).

#### MAPit software

The MAPit software was written in LabVIEW (National instruments, USA) and is based on a queued message handler, which facilitates multiple sections of code running in parallel and enables control of MAP hardware components. The main code was extended by Python-based scripts for accelerated peak detection and evaluation of a sequence of measurements.

#### Experimental modes

Generally, there are three modes for running experiments provided by the MAPit: (i) sequence mode, (ii) flow profile mode, and (iii) manual operation mode. For the first sequence mode, the user defines the pump flow rates to be measured in a sequence with an identical length of droplet generation. Additionally, the user selects the reaction temperatures to be measured from the smallest to the largest. The sequence mode enables time-resolved data collection in 12 loops on the rod, while a single loop (typically loop 12 with the longest reaction time) can be selected for end-point detection. Each data point typically consisted of at least 20 repetitions (droplets).

Second, the flow profile offers an additional level of freedom by varying the length of generation for each combination of flow rates. Practically, the flow profile mode is used exclusively with the end-point detection mode, which is measured for the entire reaction sequence and, unlike sequence mode, does not separate data based on the currently generated flow rate combination. Finally, the manual operation mode lacks the automation features of the previous two modes, yet enables quick acquisition of desired data.

#### Data acquisition modes

The data acquisition was initially performed the same way as on KinMAP.[5] To account for dispersion, a so-called “safe time-resolved” data acquisition mode was used. It works as follows: the reaction zone is first filled with the droplets corresponding to the desired combination of flow rates, and only afterwards, a signal is detected in the loops of choice in a backward direction to retrieve time-resolved fluorescence data and simultaneously prevent photobleaching. This mode was initially used for the glycerol detection pathway experiments (**Figure S8b, c**), but its efficiency, i.e., the ratio of detected to generated droplets, is relatively low (**Table S1**).

The suppressed dispersion of liquid phases mixing enabled the "step-wise end-point" data acquisition mode. This means that at each time (typically 1 min), a reaction condition is generated, and the same time is used for data acquisition. In the end, the data need to be realigned so that the particular signal matches the correct reaction condition. For this reason, at the end of each condition sequence for a particular temperature, so-called "auxiliary rows" need to be introduced to account for the conditions generated at the end of the sequence. The two modes are compared in **Table S1** and **Figure S9**.

#### Bayesian optimization scripts

A Bayesian optimization (BO) feedback loop was integrated into MAPit, providing online replanning of experiments based on the data obtained in the previous round. The BO module was written in Python and is available on GitHub: https://github.com/loschmidt/CascadeMAP-Bayesian.

Rather than using traditional random sampling, the initial experimental set is created using quasi-random Halton sequences to provide uniform coverage of the multi-dimensional parameter space for the given "budget" of conditions. The algorithm also allows combining several parameters into a simplex group, i.e., a set of parameters whose values should sum up to a fixed constant for all generated conditions. For example, joining pump flow rates of enzyme solutions in such a group enables the total flow, and thus the sum of enzyme concentrations, to be kept constant. Once a list of conditions is generated, the algorithm optimizes the execution order to minimize system stabilization delays. This is achieved via two criteria: (i) high-latency parameters (typically temperature) are changed in a single direction to avoid repeated heating and cooling cycles; (ii) for the remaining parameters, the Travelling Salesman Problem (with user-defined weights if needed) is solved to ensure the minimal overall switching distance between consecutive sets of conditions. The combination of these two criteria helped to minimize the time delays necessary for the microfluidic system to recalibrate after each experimental condition was measured.

In the implementation, the Halton series for generating the initial and subsequent sets of parameters for the limits provided by the user was implemented using the *scipy.stats.qmc.Halton* class of the SciPy library v1.17.0. The transformation of the sample for each simplex group was implemented according to the "Transformation root" method.[9] The generated conditions were sorted as follows: first, if one parameter was selected as "priority", the list was sorted according to this parameter; second, for each value of this parameter (or the entire list if no parameter is selected as priority) the weighted Travelling Salesman Problem was solved using the *solve_tsp_simulated_annealing* function of the python-tsp library.

The BO was based on the Gaussian Process Regression implemented using the scikit-learn library v1.8.0. By default, the expected improvement acquisition was used. Two different algorithms were implemented to account for batches at each iteration: Exploration-Exploitation sampling different weights in the Weighted Expected Improvement function[10] and Kriging Believer.[11] In addition to the default definitions, each one of those methods could be biased towards more exploration or more exploitation according to a separate setting available to the user. Moreover, if a selected method suggested the same conditions within one batch (which tends to happen once the method started to converge in the toy examples), our pipeline iteratively attempted to bias the current iteration to encourage more exploration for a fixed number of attempts (10 by default), after which any remaining duplicities were replaced with a random subset. The user defines the parameter ranges to be optimized and sets the "budget", i.e., the number of conditions to be tested in the initial and subsequent rounds, through a JSON configuration file (**Figure S17);** the resulting workflow is summarized in **Figures S16** and **S18**.

#### Comparison of DoE and Bayesian optimization datasets

For the comparison between the DoE optimization and the single-next-round active learning dataset of the glycerol detection pathway, 67 conditions exceeding 40 °C (out of the 292 measured BO conditions) were excluded in order to keep the same combinatorial spaces for both experiments.

For the extended multi-round runs, the reaction temperature was fixed and the "budget" was reduced from 50 conditions per round to 10 conditions in the initial round and 5 conditions per subsequent iteration. The first 48 rounds of active learning were compared, corresponding to 245 autonomously measured conditions, although for the Exploration and Exploitation (EE) algorithm a total of 100 cycles comprising 505 conditions was obtained; the further rounds were not compared, as none of the unused conditions surpassed the previously obtained maximum signal. Due to minor mechanical misalignments between the temperature-controlled incubator and the optical detection system, the raw datasets were not directly comparable across different runs. Because the initial 10-condition sequences were identical for every experiment, these points were used as an internal reference, and linear regressions were applied to project the Kriging Believer (KB) dataset onto the EE dataset (**Figure S20**).

### Reaction mixtures and off-line calibration

#### Glycerol pathway experiments

The glycerol pathway optimization experiments were usually run with the following samples in the syringes. Individual enzymes were in separate syringes at such a concentration that the final reaction concentration of all enzymes would sum to 1-3 U/mL (varying for individual experiments). The glycerol, APF, and ATP were typically mixed in one syringe, so that the final reaction mixture would contain 250 µM glycerol, 25 µM APF, and 2 mM ATP. Specific experiments required that all of these concentrations were adjusted accordingly.

The fluorophore originally used in the method of Morita and Terada,[12] Amplex Ultra-Red (AUR), gave satisfactory results on the microtiter plate reader but led to irreproducible signals and compromised droplet stability on CascadeMAP because of its precipitation in narrow capillary channels (**Figure S6**). Aminophenyl fluorescein (APF) was therefore used instead and validated on both the microtiter plate reader (**Figure S7**) and CascadeMAP.

#### TCP pathway experiments

For the TCP pathway, substrate delivery was achieved through a substrate partitioning between the oil (continuous phase) and aqueous (dispersed) phase as described previously.[5,13] The enzymes were loaded either mixed or separately in syringes with the final reaction mixture concentration of 0.6 mg/mL. The molecular weights of monomers of these enzymes (equal to their active sites) were roughly similar, so the ratios were calculated with this assumption. The reaction time was set to about 30 minutes. The solution of ~9.4 mM TCP in fluorinated oil FC-40 (3M, USA) with 0.25 % AZ2C surfactant (manufactured by CF Plus Chemicals, Czech Republic according to a previously published method[14]) was diluted at least one day ahead of the experiment by injecting TCP to the oil/surfactant mixture in a glass vial, with as little air as possible, quickly tightening and then rolling overnight on a tube roller.

For Raman analyses the concentration of TCP in oil/surfactant mixture was ~47 mM. Saturation with the substrate in the reaction mixture was required for Raman to reach limits of detection of the produced GLY. TCP was diluted in the oil/surfactant mixture in a glass vial with limited air volume by vortexing at 12,000 rpm for 120 s.

##### Off-line pH calibration

Online pH monitoring was initially attempted using a PVDF 50 Microliter flow cell (Sensorex, USA) inserted into the flowing mixture of Tricine and Bis-Tris propane (**Figure S11**). However, its large internal volume introduced significant dispersion and mixing delays, so the pH changes were calibrated off-line instead.

The dependence of pH on the buffer composition (more specifically, the Tricine percentage in the buffering system mixture) was measured in 100 μL aliquots for the glycerol detection pathway. These samples were divided to contain 4 different enzyme ratios, roughly capturing the extremes and middle of the enzyme ratio ternary plot. The pH measurements were done first for samples without glycerol (Penta, Czech Republic), ATP (Merck, USA), and APF (Merck or Thermo Fisher, USA), and subsequently after their addition (thus initiating the cascade reaction). The addition of these reaction components generally resulted in a decrease in pH, whereas the pH remained relatively constant during the reaction. Such calibration was repeated for the temperatures: 25, 30, 35, 37, and 40 °C, corresponding to the experiments on CascadeMAP (**Figure S10, Table S2**).

##### Microtiter plate reader experiments

The microtiter plate measurements were performed on the CLARIOstar plus reader (BMG Labtech, Germany) using BRAND F 96 well plates (BRAND, Germany) with 100 μL reaction volume. For measurements with AUR (Thermo Fisher, USA), the excitation and emission were set to 530-20 and 585-10 nm, respectively, while for measurements with APF, the excitation and emission were set to 485-10 and 525.5-19 nm, respectively.

#### Preparation of enzyme samples

##### Glycerol detection pathway enzymes

Glycerol kinase from Cellulomonas sp. (G6142), Glycerol 3-phosphate oxidase from Pediococcus sp. (G9637), and Horseradish peroxidase (SRE0082) were purchased from Merck, USA. Upon dissolution, the enzyme concentrations were adjusted to 40 – 100 U/mL (units were used as defined by the manufacturer) and then flash-frozen in liquid nitrogen and stored at -70 °C for further use. For CascadeMAP measurements, the enzymes were transferred to 50 mM Tris-$SO_4$ buffer (50 mM Tris-$SO_4$, 5 mM $MgSO_4$, pH 8.0).

##### TCP degradation pathway enzymes

Genes encoding DhaA31 and EchA wt were cloned into the pET21b and pET28b plasmid, respectively, in frame with a C-terminal His-tag, while the gene for HheC wt was cloned into the pET21b plasmid without any tag. The plasmids were transformed into E. coli BL21 (DE3) cells, and the proteins were expressed there. DhaA31 and EchA were purified from cell lysates by immobilized metallo-affinity chromatography using a HisTrap HP column (Cytiva, USA) charged with Ni2+ ions. HheC wt was purified using anion-exchange chromatography[15]: the crude extract was applied to a 20 ml Hi Load 16/10 (GE Healthcare, USA) packed with Q Sepharose High performance. The buffer system consisted of TEM buffer A (50 mM Tris-$SO_4$, 1 mM EDTA, 1 mM DTT, pH 7.5) and TEM buffer B (50 mM Tris-$SO_4$, 1 mM EDTA, 1 mM DTT, 0.45 M $(NH_4)_2SO_4$, pH 7.5). HheC was eluted with a two-step increasing linear gradient: 0 to 45 % buffer B in 15 column volumes (CVs) and 45 to 100 % buffer B in 3 CVs. For part of the fractions, the pure tetrameric fraction of HheC was isolated by subsequent size exclusion chromatography using a HiLoad 16/600 Superdex 200 column (Cytiva, USA).

Samples of DhaA31 and EchA wt in the 50 mM PB buffer (41 mM $K_2HPO_4$ and 9 mM $KH_2PO_4$, pH 7.5) were lyophilized for long-term storage. The HheC wt samples, both after anion exchange and after size-exclusion chromatography (in TEM B buffer or 50 mM Tris-$SO_4$, respectively), were flash-frozen in liquid nitrogen and stored at -70 °C for further use. For CascadeMAP measurements with fluorescence detection, the enzymes were transferred to 50 mM Tris-$SO_4$ buffer, while for measurements with Raman detection, the reaction buffer was 50 mM phosphate buffer.

#### Raman detection mode

##### Hardware – Raman setup

An in-house-built Raman microspectrometer was developed at the Czech Academy of Science (ISI CAS). The high-power Verdi-V6 (Coherent, USA) laser was employed to provide up to 6 W in continuous mode at a wavelength of 532 nm. The laser was passed through multiple optical components, such as mirrors and lenses (Thorlabs, USA), a 532 nm BrightLine dichroic beamsplitter (Semrock, USA), and an in-house-built anti-reflective at 540–700 nm two-way mirror for the imaging camera Watec WAT-902H (Watec Cameras, USA). To reach the sample volume, the laser was focused through an Olympus PLN 20x objective (NA 0.4) (Olympus, Japan). For proper focusing, the frame with the sample holder was mounted onto an LX30/M 25 mm XYZ Translation Stage (Thorlabs, USA). The scattered light was collected through the same objective and passed to a 532 nm StopLine single-notch filter (Semrock, USA) to pass the Raman-shifted light. An Acton SP-2556 spectrometer (Teledyne Princeton Instruments, USA) featuring a 1200 gr/mm diffraction grating, and an Andor Newton DU920P Bx-DD CCD camera (Oxford Instruments Andor, UK) were applied to separate individual wavelengths and detect their intensity (**Figure S12a**).

##### Hardware – microfluidics

Microfluidic setup was kept as similar as possible to the design used for the fluorescence assay, although some changes were necessary. Due to the high-power laser, it was not possible to analyze through the PFA tubing (ID 0.5 mm, OD 1.6 mm); therefore, a glass capillary (ID 0.5 mm, OD 0.7 mm) (both tubing and capillary from CM Scientific Ryefield, Republic of Ireland) was integrated. The capillary was held in a custom-made 3D printed holder with the same footprint as a microscope slide (26 × 76 mm) to make it universal for other standard Raman instruments. The holder was fixed to an aluminum frame (**Figure S12b**). The transition from tubing to capillary was achieved by utilizing a small piece of tubing with a larger ID (0.75 mm) that was put over the capillary as a "sleeve". The capillary with the sleeve was then fitted into the union connector JR-1066 (BGB Analytik Vertrieb, Germany), which was connected to the tubing at the other end (**Figure S12c**). This provided uniform ID throughout the system, preventing the droplets from tearing.

However, the addition of a capillary seemed to alter the behavior of the microfluidic setup, as the 6-inlet manifold P-170 (IDEX Health & Science, USA) was not efficient in forming cohesive droplets, so it was used only for precise enzyme mixing, and Y-junction P-512 (IDEX Health & Science, USA) was inserted to inject the oil into the system to create stable droplets. The total flow rate in the system was 600 μl/h. The total time of the reaction mixture (from the droplet-generating point to the detection zone) in the system was 40 min, although only 30 min was in the heated bath TVL 004 (INGOS, Czech Republic) set to 37 °C to propagate the reaction. The pumps with the syringes containing reaction solutions were kept in a Biosan Orbital Shaker-Incubator ES-20/80 (Biosan, Latvia) at 12 °C to stabilize the enzymes.

#### Data acquisition, processing, and analysis

Custom-made acquisition software based on LabVIEW (National Instruments, USA) was written for the Andor Newton DU920P Bx-DD CCD camera to automatically detect the presence of the aqueous phase in the stream of droplets and initiate Raman spectra collection. For data processing and analysis, other custom-made Python scripts were used – available on GitHub: https://github.com/pavlinasik/RAMSPECT.

To minimize unnecessary data storage, spectra were collected automatically only when the water peak intensity reached a selected threshold, which was chosen experimentally using a continuous flow of the aqueous phase (no droplets). The acquisition time was 0.1 s with 2 accumulations. This resulted in approximately 1-2 spectra collected per single droplet once they were introduced. The spectra were collected for a total of 10 minutes per enzyme ratio. From previous experiments it occurred that 2 minutes were required for the system to settle when the ratios changed. Thus, these spectra were omitted for the further processing. That resulted in approximately 750 spectra per enzyme ratio. Since the dehalogenation analysis was carried out three times and the data were averaged, the total acquisition time per enzyme ratio was about 7.5 minutes. The laser output power was set to 6 W, leading to a measuring power (the light actually coming out of the objective and into the sample) to be approximately 4.5 W.

The workflow of data processing consisted of the following steps: smoothing by Savitzky-Golay algorithm (polynomial order of 1, width 7 pixels), cropping the spectra (2700-3100 cm-1, area with all peaks of CH vibrations of the analyzed metabolites), min-max normalization of all spectra, grouping the spectra by the applied enzyme ratios, averaging within the group, and subtracting processed averaged blank signal (blank measurements were processed in the same way, the reaction mixture did not contain substrate). Then, polynomial background removal with fixed anchor points (polynomial order of 4, anchor points outside of the CH-peak area - separated by 10 cm-1 from 2700 to 2800 cm-1 and from 3050 to 3100 cm-1) was applied to decrease the residual background signal. Non-negative least squares approach (NNLS)[16] was employed to decompose the resulting spectra of the reactions and establish the contribution of TCP, DCP, and GLY Raman signals.

#### Analysis of BO datasets by agentic AI systems

The analysis of BO datasets was performed by Theorema Lab 1.0.9[17], Claude Opus 5,[18] Deepseek V4 Flash,[19] GLM 5.2[20] and Kimi K3.[21] All sessions were performed between August 25-28, 2026 using the harness engineering approach.[22] The input datasets are available on Zenodo (**Data availability**). The prompts and descriptions of the analytical methods applied by the agents are summarized in **Supplementary Analysis of CascadeMAP Experiments**. The full chat history, codes, output files and usage reports are available on Zenodo (**Data availability**). Where price per prompt could not be retrieved directly from the usage reports it was back-calculated based on the number of consumed tokens and provider price lists (**Table S4**).

##### Weighted Deming regression to check processing of peaks

For prompt 01, agreement between the peak means from automated detection script and from post-processing by the particular agentic system was manually assessed (i.e., not by the agentic AI system) using weighted Deming regression, an errors-in-variables approach appropriate when both measurement procedures are subject to random error. For each matched event, the script output and post-processed value were paired, and their respective standard uncertainties (SD) were propagated as observation-specific weights. Regression was implemented in Python using orthogonal distance regression (ODR) from SciPy, with per-observation standard deviations supplied for both axes. The fitted model yielded estimates of intercept and slope together with asymptotic 95% confidence intervals. The criterion for satisfactory results was given by the formula $|slope - 1| \leq \Delta$, where slope refers to the weighted Deming regression slope and $\Delta$ is the proportional difference of 5%, so $\Delta = 0.05$ (**Figures S21** and **S22**). All analyses were performed using a custom Python script, available on Zenodo (**Data availability**).

### Supplementary Analysis of CascadeMAP Experiments

#### Prompts

##### Note on file names in prompts

The dataset entitled "Test_exp_BO.zip" denotes the BO experiment measured with the Exploration & Exploitation algorithm, while the dataset in the folder entitled "Exp_09" corresponds to the one acquired using the Kriging Believer algorithm.

##### Prompt 1

Work with the dataset in Test_exp_BO.zip

Perform the following tasks:

1. Process and analyze the peaks in each TDMS file (channel Section Avg. vs time) from the main folder and provide a table of means and standard deviations for each of TDMS file as a separate row in the table.
2. Map the beginnings and ending of each TDMS file from the main folder onto data stored in TDMS files located in the CONT folder. For each TDMS file in CONT folder visualize the Signal (channel Section Avg. vs. time) and include vertical straight lines marking the beginnings and ends of each TDMS file from the main folder. Place also the means and standard deviations to the plot - for each TDMS file from main folder place the mean + SD as a horizontal line marking the mean and shaded rectangle denoting the standard deviation and also display the values mean + SD in the same font size as the axes tick marks. Place these in between the lines marking the start and end of particular condition.

Provide in the end information about the token usage, how many steps and sources were needed and how much time did you need to come up with the results. Perform the /usage command and save the outcomes of this to a text file usage_01.txt.

##### Prompt 2

Work with the extracted dataset from Test_exp_BO.zip

Perform the following tasks:

1. Align the pump flow rates with the corresponding means and SDs (taking into account only non-outlier droplets for each TDMS file) following the logic of **Data collection** described bellow. Create a csv file with only the main conditions and their relevant means and SDs from all the aligned conditions. Take into account that there were always 5 auxiliary conditions per Bayesian optimization (BO) round. Initial BO round consisted of 10 main conditions, while all following BO rounds contained 5 main conditions per round.
2. Plot a graph showing the dependence of cumulative maximum signal per BO round (y axis) over consecutive BO rounds.

Provide in the end information about the token usage, how many steps and sources were needed and how much time did you need to come up with the results. Perform the /usage command and save the outcomes of this to a text file usage_02.txt.

**Data collection**

The data collection for each experiment proceeded as follows:

1. The initial set of conditions (initial round) was scheduled based on the number of conditions and the parameter ranges (in this case, just the limits of a particular pump's flow rates).
2. Within a particular round, each set of conditions was augmented by putting 5 auxiliary conditions at the end to account for what we call step-wise endpoint detection. This means that we simultaneously record the data (each condition in a separate row, i.e., one file) and generate droplets for a particular condition. For example, if we set the (data) acquisition time of 60s, then from time 0s to 60s we have generated the first condition in the schedule and at the same time recorded row no.1. However, the data in row no.1 don't correspond to condition no.1, because at that time the droplets for row 1 had not yet reached the endpoint detection. We need to take row no.6 to learn about data corresponding to condition no.1. This is given by the reaction time of approximately 5 minutes, while the data acquisition time is a fraction of this reaction - actually, it is divided by a whole number. Therefore, after all data for the main and auxiliary rows (conditions) have been acquired, the signals evaluated for each row by the peak-processing script need to be aligned with specific conditions (flow-rate combinations).

#### Prompt 3

Work with the csv table of main conditions from the previous step.

Perform the following tasks:

1. Perform a principal component analysis (PCA) including the parameters of HRP, GPO, GK, BTP, Tricine and means of signal. Generate a score plot of first two principal components including the % of data it explains.
2. Perform the SHapley Additive exPlanations (SHAP) analysis and generate the beeswarm plot reflecting the impact of HRP, GPO, GK, BTP and Tricine on the mean signal.
3. Evaluate the degree of exploration and exploitation and plot these values per round in dependence of consecutive rounds.
4. Think of other types of multi-parametric analysis and visualizations which might give us insights to the Bayesian optimization experiment.

Provide in the end information about the token usage, estimated price in $, how many steps and sources were needed and how much time did you need to come up with the results. Perform the /usage command and save the outcomes of this to a text file usage_03.txt.

#### Prompt 4

Work with all the results from previous steps.

Produce an interactive html file with the following contents:

1. General information about the experiment, including names of individual parameters and number of main and auxiliary conditions.
2. Quality control panel: The graph of mean signal per each main condition (y axis) and Bayesian optimization (BO) round on the X axis. By clicking on a particular datapoint a corresponding plot from continuous plots similar to the outcome 2 from prompt_01.md should appear with the highlighted condition.
3. Interactive graph of cumulative maximum signal and maximum signal per round (both on Y axis) shown in dependence of consecutive BO rounds - similar to the outcome 2 of prompt_02.md.
4. Interactive graphs of the outcomes of prompt_03.md, including PCA score plot, SHAP beeswarm plot, progress of exploration and exploitation throughout the BO experiment and other analyses which came out of the prompt_03.md.

Provide in the end information about the token usage, estimated price in $, how many steps and sources were needed and how much time did you need to come up with the results. Perform the /usage command and save the outcomes of this to a text file usage_04.txt.

#### Prompt 5

Work with the previous dataset Test_exp_BO and with a new dataset from folder Exp_09. Dataset Exp_09 has the same structure as Test_exp_BO.

Perform the following tasks:

1. Run the prompts from files prompt_01.md, prompt_02.md, prompt_03.md and prompt_04.md but this time for Exp_09 dataset, not with Test_exp_BO.
2. Merge the outputs of prompt_04.md - the interactive HTML file - to a single HTML file, which would include both Test_exp_BO and Exp_09. Here are some further desired details about the final HTML file:
    - 2.1. Rename the datasets so that Test_exp_BO = Exploration & Exploitation and Exp_09 = Kriging Believer.
    - 2.2. Convert the flowrates of individual solutions to the percentages, so that for enzyme solution HRP, GPO and GK, 6 ul/min = 100 % and for buffers Tricine and BTP, 4 ul/min = 100 %.
    - 2.3. For every panel/graph include a short description (max. 1 sentence) so that the meaning/data can be understood by non-experts.
    - 2.4. Ensure that there are no graphical mistakes, such as overlapping graphs, text not properly fitting the boxes and others.
    - 2.5. For every graph the user should be able to filter the particular dataset.

Provide in the end information about the token usage, estimated price in $, how many steps and sources were needed and how much time did you need to come up with the results. Perform the /usage command and save the outcomes of this to a text file usage_05.txt.

#### Analytical methods

##### Claude Opus 5

**Datasets and data acquisition**

Two Bayesian-optimisation campaigns of a three-enzyme cascade — glycerol kinase (GK), glycerol-3-phosphate oxidase (GPO) and horseradish peroxidase (HRP), with a peroxidase-coupled optical readout — were re-analysed. Reagents were delivered by seven syringe pumps into a segmented (droplet) flow and interrogated by a CCD detector; the analysed quantity throughout is the Measured Data / Section Avg. vs. Time channel, an optical intensity in arbitrary detector counts, sampled at 20.05 Hz (50 ms CCD integration). All acquisition parameters below are as recorded in the instrument metadata. Campaign EE comprised 1005 acquisition rows and 67 continuous-recording files over 18.5 h (2025-02-11/12); campaign KB comprised 485 rows and 33 continuous files over 8.9 h (2025-02-13). Each row covered 57.5 s (median). Five streams were varied by the optimiser (HRP, GPO, GK, and the buffers BTP and Tricine); the mixed reagent stream (2.0 µL/min in EE, 1.0 µL/min in KB), the segmenting carrier (12.0 µL/min) and temperature (25 °C) were held constant. Because the enzyme streams sum to 6.0 µL/min and the buffers to 4.0 µL/min, flow rates are reported throughout as a percentage of the corresponding group total, each stream spanning 10–80 % (enzymes) or 15–85 % (buffers). Each optimisation round scheduled its main conditions and appended five auxiliary conditions of one fixed flush composition (HRP, GPO, GK = 1.0; BTP, Tricine = 4.0 µL/min), identical in both campaigns. EE ran 100 rounds (10 + 99 × 5 = 505 main conditions); KB ran 48 of 50 planned rounds (10 + 47 × 5 = 245).

**Droplet detection and per-condition statistics**

Within each acquisition record, droplets were detected as contiguous excursions above a threshold placed 10 % of the way from the 5th to the 99.5th percentile of that record — deliberately close to the carrier baseline rather than at half height, so that droplets of widely differing intensity within one record are all detected. Excursions shorter than eight samples (0.4 s) were discarded, as were excursions truncated by the record boundary. The intensity of a droplet was taken as the mean of its contiguous flat top, defined as the samples within the upper 10 % of that droplet's own amplitude, which excludes the rising and falling flanks. Outlying droplets were removed by a median/MAD rule ($|x - \text{median}| > 3.5 \times 1.4826 \times \text{MAD}$), applied per record; this is robust to up to 50 % contamination and makes no assumption about where in the record the outliers occur, which matters because both carry-over at condition boundaries and isolated bubbles occur. This removed 743 of 21 617 droplets (3.44 %) in EE and 636 of 20 162 (3.15 %) in KB, at most six per record. Each condition is summarised by the mean and standard deviation of its retained droplet intensities (median 21 droplets per condition in EE, 42 in KB; median within-condition CV 3.0 % and 5.5 %).

**Placement on the continuous recording and condition–signal alignment**

The per-record intensity arrays proved to be verbatim sub-arrays of the concatenated continuous recording, so each record was located by exact array matching (search on the first sample value through a sorted index, followed by full-array verification), requiring no clock-drift or offset model. Absolute timing was taken from the CCD OcO Timestamp channel (LabVIEW epoch, seconds since 1904-01-01 UTC; displayed as CET). Because reagents require approximately five minutes to reach the detector while each record spans 57.5 s, the droplets of the condition being pumped during row n are recorded in row n + 5; the five auxiliary conditions per round exist to flush the last main conditions of that round through to the detector. This five-row offset was verified in four independent ways: the pump settings stored in each record's metadata reproduce the corresponding row of the measured flow-rate log exactly (maximum deviation $5 \times 10^{-5}$, versus 3.87 at offsets of ±1); the main-condition rows of that log reproduce the optimiser's own schedule file; the correlation between droplet intensity and the auxiliary indicator inverts sign between lag 0 and lag 5, with its extremum at exactly 5; and the auxiliary-condition response is flat across all five positions of the auxiliary block, which an off-by-one offset would destroy. All 505 EE and 245 KB main conditions received a signal; the only conditions without one are the five auxiliary conditions of the final round, as the design predicts.

**Multivariate analysis**

Principal-component analysis was performed on the five flow rates plus the mean condition intensity, z-scored (correlation-matrix PCA) and computed by singular-value decomposition. Feature attribution used exact Shapley values: with five features the $2^5 = 32$ coalitions were enumerated in full, using the interventional (marginal) value function with the complete set of observed conditions as background, so no sampling or kernel approximation was involved. The implementation was validated against analytic cases to $\sim 10^{-16}$ (additive decomposition, efficiency, symmetry and null-player) and satisfied efficiency to $7 \times 10^{-12}$ on the fitted model. Attribution requires a model; we used a random forest (300 trees, depth 12, three features per split) whose predictive accuracy was quantified by five-fold cross-validation ($R^2$ = 0.932 in EE, 0.568 in KB) and corroborated by permutation importance. Because BTP and Tricine are exactly anti-correlated, their contributions are not separately identifiable; the whole attribution was therefore repeated on the three free coordinates (HRP, GPO, BTP), where every evaluated point remains within the feasible composition set, and gave the same ordering. Pairwise interaction strength was quantified as the non-additive fraction of a two-way partial-dependence surface (a Friedman-style $H^2$ statistic).

**Search-behaviour metrics**

The acquisition function itself is not recorded in the data, so exploration and exploitation were quantified geometrically in the three-dimensional feasible space (HRP, GPO, BTP scaled to their design bounds). For each round we computed, for every proposal, the distance to its nearest already-evaluated condition (novelty) and the distance to the current best condition (proximity), and normalised both by the median nearest-neighbour spacing of the evaluated history. This normalisation is necessary because raw nearest-neighbour distances shrink as n^(−1/3) as the space fills, which would otherwise be mistaken for a decline in exploration. Individual proposals were classified as exploitative when within one spacing of the incumbent, exploratory when further from all prior conditions than that spacing, and intermediate otherwise. These are behavioural descriptors of where the optimiser sampled, not statements about its internal acquisition criterion.

**Reproducibility assessment**

Three independent estimates of measurement reproducibility were compared. First, the fixed flush composition, being identical in every round, provides genuine replicates spread over the whole campaign ($n = 99$ in EE, $n = 47$ in KB); its between-round standard deviation is the primary estimate. Second, an empirical semivariogram of condition intensity against composition distance, together with the intensity difference between near-duplicate condition pairs, provides an upper bound on the same quantity. Third, the within-condition standard error of the mean over droplets quantifies only droplet-to-droplet scatter within a single 57.5 s window. A global exponential variogram model did not identify a nugget here — the semivariance rises out to the largest separations — so the replicate-based estimate is reported in preference to a fitted nugget. Stability of the flush response over time was tested by rank correlation against round index ($\rho = -0.109$, −0.07 % per round in EE, consistent with no drift; $\rho = -0.281$, −0.39 % per round in KB, a weak negative trend of the same order as the round-to-round scatter). Since the flush composition is identical in both campaigns while the mixed reagent stream is not, intensities are additionally reported normalised to the concurrent flush response wherever the two campaigns are compared.

**Software and reproducibility**

Analysis was performed in Python 3.11.15 with numpy 2.4.3, pandas 3.0.5, scipy 1.17.1, matplotlib 3.11.1 and npTDMS 1.11.0 for reading the instrument files. Because domain-specific packages were unavailable in the analysis environment, the principal-component analysis, the random-forest surrogate with out-of-bag and cross-validated scoring, permutation importance, the exact Shapley enumeration and all interactive figures were implemented directly on numpy primitives; each implementation was validated against analytic or independently computed references before use, as described above. The analysis pipeline is parameterised by dataset, so both campaigns pass through identical code paths; re-running the first campaign after the second was added reproduced its results exactly. Condition-level tables, per-round search metrics, attribution values and the complete set of figures are provided as supplementary files, together with a self-contained interactive HTML application that reproduces every figure for either campaign or both.

#### Deepseek V4 Flash

**Data parsing and time-base restoration**

TDMS files (one per recording row, plus contiguous CONT segments) were read with nptdms. In every file the Section Avg. vs. Time signal is logged at 20 Hz whereas the internal Time (s) channel is recorded at a lower rate (≈1 Hz in row files; ≈10 Hz in CONT files), causing naive x/y plotting to fail. A uniform 20 Hz time axis was therefore reconstructed for each signal by linearly interpolating the Time (s) channel onto the signal sample indices (numpy.interp over index space), giving a monotonic absolute-time axis for every trace. For each detector row, the steady-state flow rate of each channel was taken as the median of the Pump1–Pump7 channels.

**Peak processing and outlier cleaning**

The Section Avg. vs. Time trace of each row is a train of droplet spikes superimposed on a low baseline (~250 a.u.), so a single scalar per row was not taken from the raw waveform (whose mean lands mid-peak) but from its detected peaks. Droplet peaks were located with scipy.signal.find_peaks using a minimum inter-peak distance of 25 samples and a height threshold set adaptively to the midpoint between the robust low and high modes, $h = (P5 + P95)/2$, where $Pq$ is the q-th percentile of the signal. For each row the peak heights were collected and outlying droplets discarded as heights lying more than k·1.4826·MAD from the median height (MAD = median absolute deviation, k=3); the retained peak heights defined that row's condition statistic, reported as mean ± SD. This yields a statistic localized to the droplet-peak plateau rather than diluted by baseline excursions.

**Condition alignment**

Because a droplet generated in row j reaches endpoint detection only after the ~5 min reaction time (five 60 s rows), the signal recorded in row j was attributed to the pump conditions set in row j−5, i.e. condition c ↔ pump row c and signal row c+5. Rounds are structured with the initial round containing ten main conditions and all subsequent rounds five main conditions, each round augmented by five auxiliary conditions for stepwise endpoint detection; the number of main conditions per experiment was taken directly from the logged set-point table (all_sorted.csv, 505 and 245 rows for the two experiments) and the condition-to-row placement was derived under this 10/5+5 scheme, then validated by matching each set-point vector to the measured per-row flow-rate log (25.00degC_FlowRates-Measured-log.csv) within a value tolerance of $\leq 1\times10^{-2}$ across HRP, GPO, GK, BTP and Tricine; for Exp_09 the flow-rate log's first column is a flag (unity) rather than the MIX flow, and the comparison was accordingly restricted to the five tunable channels. The final aligned table contained, per main condition, the BO round, condition indices, pump-row and signal-row pointers, the five pump flow rates, and the aligned cleaned peak mean and SD.

**Multivariate analyses**

Principal-component analysis (PCA, sklearn.decomposition) was applied to the standardized (z-scored) matrix of the five pump rates plus mean signal, and the projection onto the first two components (accounting for 34.0% and 33.3% of the variance in the primary run) was plotted as a score plot with loading vectors. Feature attribution used Tree SHAP (shap.TreeExplainer) on a gradient-boosted regression surrogate (scikit-learn GradientBoostingRegressor, 300 estimators, maximum depth 3, learning rate 0.05, random_state=0) that predicted mean signal from the five pump rates; results were reported as beeswarm plots (SHAP value colored by feature value) and mean-|SHAP| summaries. Partial-dependence curves were computed with sklearn.inspection.partial_dependence (30 grid points per feature), and pairwise Spearman correlations (5×5 channel matrix plus signal) were computed over all conditions.

**Exploration/exploitation and convergence**

For each BO round, the five proposed recipes were normalized to the [0,1] per-channel range across the whole experiment. Exploration was the mean, over a round's recipes, of the normalized Euclidean distance to the nearest previously sampled recipe; exploitation was the mean of 1 − min(1, d_best/D_max), where d_best is the distance to the current best-so-far recipe and D_max the maximum pairwise distance observed. Round maxima and the running cumulative maximum signal (normalized by D_max) tracked convergence.

**Interactive consolidated dashboard**

Per-condition statistics, downsampled continuous traces (≤1,200 points per CONT segment via uniform decimation), PCA scores/loadings, SHAP values, exploration/exploitation series and partial-dependence data were serialized to JSON and embedded into a single self-contained HTML file using Plotly (Plotly.js served locally for offline use). The dashboard renders a quality-control scatter of mean signal versus BO round in which selecting a point loads that condition's continuous Section Avg. vs. Time trace with its mean ± SD band and start/end boundaries; it also includes the cumulative-maximum series and all multivariate plots, each with a per-dataset filter (Exploration & Exploitation vs. Kriging Believer) and a one-sentence plain-language caption. Flow rates were displayed as percentages of full scale (HRP/GPO/GK, 6 µl/min = 100%; BTP/Tricine, 4 µl/min = 100%). Rendering, filtering and click interactions were verified headlessly in Chromium via Playwright (zero JavaScript errors; correct trace counts per filter state).

**Software and compute environment**

All analyses were performed in Python 3.11.15 on Windows 11, using NumPy, pandas 3.0.5, SciPy 1.17.1, scikit-learn 1.9.0, SHAP 0.51.0, Matplotlib 3.11.1, nptdms 1.11.0, openpyxl 3.1.5, and Plotly.js 2.32.0; interactive-dashboard verification was performed headlessly with Playwright (Chromium). Deterministic training was fixed with random_state=0. Computations were run on a standard workstation; the full pipeline (parsing, peak analysis, alignment, and all multivariate analyses for both experiments) completed in minutes.

#### GLM 5.2

Continuous fluorescence readouts were recorded as TDMS files and processed in Python 3.11 using the nptdms (v1.11.0), NumPy (v2.4.3), SciPy (v1.17.1), pandas (v3.0.5), and matplotlib (v3.11.1) libraries. For each experimental condition (n = 1,005 row-based TDMS files), the mean spectroscopic-section signal ("Section Avg. vs. Time" channel, ~1,150 points per file) was extracted as the analytical readout. Because each recording exhibited a strongly bimodal intensity distribution — arising from periodic droplet transitions between aqueous plugs, which produce a low-intensity baseline (~250–1,000 a.u.), and the fluorescence signal of interest, which forms a high-intensity plateau — the peak amplitude was isolated by unsupervised two-class k-means clustering (k-means++ initialization, fixed seed for reproducibility) of all intensity values within a file; points assigned to the higher-centroid cluster were retained as the peak signal. To further reject intermediate values arising from transitions between the two intensity states, an interquartile-range filter (1.5×IQR fence applied to the retained cluster) was applied, and the peak height and its variability were quantified as the arithmetic mean and sample standard deviation of the remaining points (~52% of the raw record per file on average). To contextualize the discrete, condition-based measurements within the uninterrupted experimental timeline, the start and end times of each row-based acquisition were mapped onto the continuously recorded signal (67 segmented TDMS files, ~1,000 s each), using the CCD acquisition timestamps aligned to the experiment's internal clock. For each continuous segment, the raw trace was visualized together with the boundaries of every overlapping condition, and the corresponding cleaned peak statistics were annotated as horizontal lines (mean) and shaded rectangles (mean ± SD) within each condition window, enabling direct visual assessment of peak stability and drift across the experiment.

#### Kimi K3

Raw data were processed offline in Python 3.11 using the nptdms (v1.11.0), NumPy, pandas and matplotlib libraries. For each condition file, the waveform channel "Section Avg. vs. Time" (1 Hz sampling) of the group Measured Data was extracted together with its absolute acquisition start time (wf_start_time). Individual reaction peaks were localized by first determining the global signal maximum and the local post-peak baseline (mean of the terminal five samples; for truncated peaks the pre-peak minimum was used instead), then expanding the peak window symmetrically from the maximum until the signal fell below 10 % (leading edge) or 15 % (trailing edge) of the peak height above baseline. To ensure that the resulting statistics described the peak height rather than the mean of the concentration ramps, each window was further trimmed to a flat plateau by requiring boundary samples to lie within 10 % of the peak height above baseline, and outlying "droplet" samples within the plateau were removed using a robust two-sided median-absolute-deviation (MAD) filter (rejection at 3.5 × 1.4826 × MAD around the sample median). Peak means and standard deviations were computed from the cleaned plateau samples only (sample SD, ddof = 1); 45 files whose peaks extended beyond a recording boundary (maximum at the first or last sample) were processed using their observable top samples against a pre-peak-minimum baseline and were explicitly flagged as truncated. For temporal context, the absolute plateau windows (acquisition start time plus sample offsets) of all condition files were intersected with the time spans of the 67 continuous recordings, and the continuous traces were visualized with vertical markers at each condition's onset and termination, overlaid with the corresponding plateau mean (horizontal line) and mean ± SD (shaded band) and annotated with the numerical values in collision-checked, staggered label lanes. All 1 005 condition peaks were successfully quantified; in total 2 788 ramp/droplet samples were excluded from the statistics, reducing the median within-peak standard deviation from ~2 441 to ~52 counts, i.e. the reported means robustly represent plateau-level signal intensities.

#### Theorema Lab 1.0.9

##### Droplet-resolved signal extraction

Raw fluorescence traces were read from TDMS acquisition files with the `nptdms` library, using the "Section Avg. vs time" channel and the internal experiment clock ("Time (s)") of the "Measured Data" group. Each trace is a droplet train — a low inter-droplet baseline punctuated by discrete droplet plateaux — so summarising a trace by its overall mean is inappropriate. Individual droplets were segmented per file by Otsu thresholding of the intensity distribution; connected above-threshold regions separated by gaps of ≤ 5 samples were merged, and merged regions spanning ≥ 10 samples were retained as droplets. Each retained droplet was reduced to the mean of its samples. Droplet-level outliers were then removed within each acquisition window using a robust criterion, median ± 3 × median absolute deviation, and the objective for that window was taken as the mean of the surviving droplet means, with the corresponding standard deviation reported as within-condition dispersion. Across the E&E campaign this retained 20,098 droplets and excluded 2,021 (9.1%). The adequacy of the segmentation was checked by comparing the coefficient of variation of droplet-resolved means against that of whole-trace means (**Results**); a droplet-resolved CV approaching the whole-trace value would indicate that baseline had leaked into the retained regions.

##### Continuous-recording cross-check

Alongside the per-condition acquisition windows, the experiment was recorded continuously and split into files of approximately 1000 s (67 files in E&E, 33 in KB). Each main acquisition window was mapped onto the continuous recording via the relationship between the protocol timestamp of the "Protocol Status" group and the CCD timestamp channel of "Measured Data", and every continuous file was rendered with vertical markers at the start and end of each overlapping window and the corresponding droplet-resolved mean ± SD annotated between them. This provides a per-condition visual audit of segmentation and alignment against the raw signal.

##### Multivariate analysis

Principal-component analysis was performed twice on each campaign: once on the five standardised flow rates together with the objective, as a direct description of the delivered design, and once on an isometric log-ratio (ILR) parameterisation of the two sub-compositions together with the objective, which removes the rank deficiency imposed by the closure constraints. Explained-variance ratios are reported for both; components of the first parameterisation carrying variance at the level of numerical noise are identified as constraint artefacts rather than as structure.

A random-forest regressor was fitted to predict the droplet-resolved objective from the five flow rates, and TreeSHAP attributions were computed on the fitted model. Model adequacy was assessed both by 5-fold cross-validation and, because the data are a time-ordered optimisation trajectory rather than an i.i.d. sample, by a chronological hold-out in which the model was trained on early rounds and tested on the final rounds; both are reported, and attributions are interpreted only in the light of them. Since the members of each fixed-sum stream are mutually determined, per-feature Shapley values are not individually identifiable; attributions are therefore additionally reported as coalitions, {HRP, GPO, GK} and {BTP, Tricine}, and conclusions are drawn at the coalition level only.

##### Exploration and exploitation metrics

Sampling behaviour was quantified per round in the standardised, constraint-aware coordinate system. Exploration was defined as the mean distance from each condition of a round to its nearest already-sampled condition from any earlier round; exploitation was defined as the mean proximity of the round's conditions to the incumbent best condition at the start of that round. Within-round diversity was recorded as the mean pairwise distance among a round's own conditions. Reporting all three separates two behaviours that a single metric conflates: sampling that contracts around the incumbent, and sampling that contracts around a region that does not contain it.

##### Reproducibility

Analyses were executed through an automated analysis backend under version control of the input datasets; every derived table and figure referenced here is retained with the identifier of the run that produced it. Condition-level values were spot-checked against the source acquisition tables, and both campaign schedules were reconstructed from their per-round files and compared with the campaign-level schedule before use. An interactive supplementary file provides the per-condition quality-control view, the optimisation trajectories, and the multivariate analyses for both campaigns, with the raw continuous traces embedded so that any individual condition can be inspected against its underlying signal.

17. Theorema. Theorema — Autonomous R&D Agents. https://theorema.ai.

18. Introducing Claude Opus 5. https://www.anthropic.com/news/claude-opus-5.

19. DeepSeek-V4 Preview: Entering the Era of Affordable Million-Token Context. https://www.deepseek.com/en/news/v4-preview/ (2026).

20. GLM-5.2: Built for Long-Horizon Tasks. https://z.ai/blog/glm-5.2.

21. Kimi K3. *Kimi API Platform* https://platform.kimi.ai/docs/guide/kimi-k3-quickstart.

22. Galster, M. *et al.* Configuring Agentic AI Coding Tools: An Exploratory Study. in *Proceedings of the 3rd ACM International Conference on AI-Powered Software* 11–20 (Association for Computing Machinery, New York, NY, USA, 2026). doi:10.1145/3805760.3814887.
